# Morphometrical asymmetries and tractography of speech-relevant cortex in relation to language lateralisation and rapid temporal processing

**DOI:** 10.1101/2024.04.03.587995

**Authors:** Jesse D Bourke, Gavin Cooper, Birte U Forstmann, Ulrich Schall, Juanita Todd

**Author notes:** Corresponding author (JDB).

## Abstract

Differences in the functional roles of the left and right cortices for speech-related processes have been known since the findings of Broca [1] and Wernicke [2]. Nearly 100 years later anatomical asymmetries of speech-related cortex was emphasised as a potential substrate to such functional lateralisations [3]. Exploration of associations of anatomical asymmetries and functional lateralisations in speech has since continued, with developing technologies and theoretical insights mutually affording increasingly refined understandings. The present study is another such continuance; we outline and report associations of neuroanatomical (morphometrical) and connective (diffusion tractography) measures of speech-related cortex with differences of participant speech lateralisation and rapid temporal acuity (a hypothesised general auditory ability that contributes to superior speech processing). Review and support of developments in methodological approaches to morphometry and tractography to are also provided. Overall, our study affirms complex and selectively overlapping relationships of anatomy and connectivity (especially in the planum temporale) with behavioural language lateralisation and the processing of rapid temporal acoustics. Implications, limitations, and recommendations are discussed.

## Introduction

Variance in the functional roles of the left and right cortices for speech-related processes have been known since the findings of Broca [1] and Wernicke [2]. Nearly 100 years later anatomical asymmetries of speech-related cortex were emphasised as a potential substrate to such functional lateralisations [3]. It is now well established that there are indeed functional lateralisations of speech-related processes [4–7] as well as structural asymmetries of speech-related cortex in temporal and frontal regions and the white matter connectivity between key regions of interest (ROI) [8, 9].

A prominent perspective of functional lateralisation is that effective ‘division-of-labour’ between the hemispheres can produce superior behavioural, cognitive, and perceptual ability [10–12]. Accordingly, speech processing is in general considered to involve processes that whilst bilateral, are differentially right-hemisphere biased or left-hemisphere biased depending on the particular variable of interest and its measurement [13]. Overall, the left-hemisphere is considered typically dominant in speech perception. There has been debate as to whether it is a biased functional role for linguistic or acoustic factors of speech driving this leftward bias. Regarding the acoustic perspective, leftward lateralisation of rapid temporal cues (and conversely rightward asymmetry of slower temporal cues) has been emphasised [14; see also ‘asymmetric sampling in time’ hypothesis: 15, 16, 17]. However, this is likely a false dichotomy and a more useful conceptualisation is that both linguistic and acoustic (especially rapid-temporal) features of speech are dynamically involved in the general left-hemisphere speech bias [for review see 18].

Following Geschwind and Levitsky [3], structural asymmetries continue to be suggested as a substrate for functional laterality and subsequent behavioural ability advantages, both in terms of macrostructure of cortex [19, 20] and their connecting white matter tracts [21–23]. However, integrative studies addressing structural-behavioural links are rare, particularly for when simultaneously examining grey and white matter measures, as well as exploring language lateralisation and associated basic acoustic auditory processes. Empirical studies conjunctively assessing structural asymmetries with functional lateralisations and behavioural abilities for speech-related processing are needed for further developments in consolidating understandings in this area.

### Study Aims

In the present study, we explored associations of morphometrical and connective (diffusion-weighted) measures speech-related cortex with behavioural speech lateralisation and rapid temporal acuity performance. Here, *morphometrical* and *connective* both refer to *structural* measures: physical properties of the brain obtained with *magnetic resonance imaging* (MRI). The planum temporale is the emphasised due to being the foremost region-of-interest (ROI) in studies of lateralisation of speech [24–27]. Surface area is emphasised as the primary variable of interest to likewise adhere to extant trends in the literature [28, 29]. Other regions (*Heschl’s gyrus* (HG), *inferior frontal gyrus pars opercularis* (IFGop), & *inferior frontal gyrus pars triangularis* (IFGtr)) and variables (volume, thickness, & gyrification) are reported in supplementary material. Connective measures of white matter were obtained using *diffusion-weighted MRI* (dMRI), specifically a method termed probabilistic tractography. *Interhemispheric* connections were measured between left to right PT across the corpus callosum. *Intrahemispheric* connections were also measured to identify asymmetry of part of the white matter bundle named the *arcuate fasciculus* (AF) that connects the PT to the IFGop.

Also reported in relation to our structural measures are individual variables of participant non-forced *dichotic listening task* right ear advantage (NF-DLT REA) performance as a measure of *language lateralisation* and *gap detection thresholds* (GDTs) as a measure of *rapid temporal processing* (RTP). Participant age, sex, and total brain volume were included as control variables.

There has been a rapid acceleration of developing technology for structural paradigms. As such, we provide a detailed methodology rationale for our structural measures both in terms of the choice of structures measured (ROI & tracts) and the variables of their measurement. Experimental evidence further affirming this rationale is then presented in supplementary material, with an aim to increase awareness and consideration to these factors in future studies. Given the primary aims pertain to structural-functional relationships, we first provide relevance and rationale of our behavioural performance measures.

### Methodology Rationale

#### Language lateralisation: Dichotic listening task

The *dichotic listening task* (DLT) is a prominent example of leftward speech lateralisation from a perspective of both linguistic and acoustic features [6]. There is a robust finding that presenting verbal stimuli (e.g., words, or consonant vowel syllables, CVSs) to the left and right-ear dichotically (e.g., /*ba*/ to the left-ear, /*pa*/ to the right-ear) produces a bias where individuals preferentially report the right-ear stimulus [30]. Given that the peripheral to central ascending auditory pathways are contralaterally dominant, this right-ear advantage (REA) is suggested to be reflective of a left-hemisphere advantage [31]. The linguistic relevance of the REA is affirmed by high (up to ∼90%) reliability ratings of the DLT with functional MRI and sodium amytal (alternatingly anaesthetising each hemisphere during a language task) based language lateralisation measures [e.g., 32, 33-35].

#### Rapid temporal processing ability: Gap detection thresholds task

Gap detection threshold (GDT) paradigms involve determining the minimally detectable “gap” between two sounds; the shorter the gap, the indicated superior RTP ability of a participant [36]. In the context of speech, the gap between a consonant (e.g., /b/) and vowel (e.g., /a/) of a consonant-vowel-syllable (e.g., /*ba*/) is referred to as *voice-onset-time* and is an established feature for categorical detection and representation of such stimuli [37–39]. For example, /*ba*/ and /*pa*/ are phonetically similar yet respectively discriminable by their shorter and longer voice onset time (∼5ms vs. ∼25-50ms), having implications for higher-order speech processes [e.g., words: “bath” vs. “path”, to sentences: “Find a [b/p]ath”, etc.; 40]. Non-speech analogues of consonant vowel syllables more directly capture acoustic RTP by using leading and trailing markers (stimuli with similar acoustic properties but no linguistic content) in place of consonants and vowels. Comprising these markers with different frequencies (i.e., stimuli selective for different auditory channels) creates a *between-channel* gap, which has been shown to be a more effective analogue of voice-onset-time. Indeed, there is supporting evidence that acoustic GDTs can (partially) account for speech detection abilities and pathologies [41–43].

Acoustic relevance of the REA in dichotic listening has been attributed to leftward lateralisation of RTP [14] following evidence of superior right-ear/left-hemisphere RTP demonstrated in GDT paradigms [44, 45]. Specifically, it has been shown that improving or reducing the clarity of RTP cues (i.e., voice-onset-time) of consonant-vowel-syllable stimuli can potentiate or attenuate the REA respectively [46–48]. Furthermore, in an event-related potential paradigm (mismatch negativity), Todd et al. [49] demonstrated an indirect (right-ear) leftward bias for non-speech consonant-vowel-syllable analogues – an effect contingent upon acuity in a behavioural GDT task (i.e., in individuals with superior RTP ability). We are unaware of pre-existing studies directly exploring behavioural NF-DLT and GDT measures. As such, the current study employs the tasks as described by Hugdahl, Westerhausen [31] and Todd et al. [49] respectively to further asses their mutual relationships as behavioural measures, as well as explore associations with structural measures.

#### Morphometrical variable and region-of-interest

The classical models [1–3] offered revolutionary early insights into the role of structural asymmetry in speech-related processes. However, with the evolution of the field it has been argued that the terms Broca’s area and Wernicke’s area are due for extinction as they misrepresent important subtleties of speech-related cortex and its morphology [50–52]. Indeed, modern neuroanatomical atlases provide remarkable advancement in consistency for measuring such structures [e.g., 53, 54, 55]. Thus, we have focused on identifying contemporary morphometrical structure parcellations that may best capture the role of asymmetries in speech-related processes. The PT, HG, and pars opercularis and pars triangularis of the IFG are viable ROI to more granularly represent the classical “Broca’s area” and “Wernicke’s area” constructs [50, 56]. We clarify the selection of the PT from these regions below, but first give context of surface area as a useful variable of interest.

#### Use of cortical surface-area

Formative studies of structural asymmetries in speech related cortex used volume as the primary variable. However, developing evidence has since shown dissociations of surface-area and thickness, with the former better explaining volumetric asymmetries [28, 29, 57–60]. Gyrification has also been acknowledged to offer further insight [29, 58], particularly given its implicated influence on surface-area and morphometrical patterns [61, 62]. Indeed, gyrification in speech-related cortex may be a factor of differences in phonetic categorisation ability [63]. However, empirical understanding of the relationship of asymmetry of this structural variable with performance and functional lateralisation of speech-related processes is not yet developed. As with most of these studies, the software FreeSurfer [www.surfer.nmr.mgh.harvard.edu; 64] was chosen for morphometrical analysis. We used the Destrieux, Fischl [53] atlas to maximize consistency in comparison with these studies [the alternative Desikan et al. atlas does not include the PT; 65].

#### Emphasis on planum temporale and surface-area

The PT is the most extensively researched region in terms of structural asymmetries, behavioural and psychophysiological lateralisations, and performance abilities in speech-related processes [66]. Although structural asymmetry has been strongly implicated as a core substrate of behavioural and functional factors [8], studies with direct linking are however limited. Indeed, behavioural ability and functional lateralisation have been shown to relate with left PT surface-area [67] and hearing acuity impairment has been linked with cortical thinness in the left HG [68]. Regarding asymmetry, cortical thickness has also been related to functional laterality and behavioural ability, though in respects to longer-temporal window processing [i.e., AST-related rightward lateralisation; 69]. At the time of study development and conduct this was (to our knowledge) the extent of literature addressing structural-functional-behavioural links of asymmetry and lateralisation in speech-related processes.

Leftward surface-area asymmetry of HG is now well established [9]. There is also some support for a link of leftward HG volume asymmetry with lateralisation and behavioural ability [70]. However, the evidence base is considerably weaker than for the PT and lacks the specific analysis of surface-area and thickness. There is good evidence of leftward asymmetry in IFGop in terms of surface-area [9] and involvement in phonological speech-related processes [71, 72]. However, a relationship between these factors has not been established to our knowledge. Furthermore, although the IFGop is often combined with IFGtr to form Broca’s area, there is evidence challenging the existence of a general leftward asymmetry of IFGtr and involvement in phonetic processing [28, 73]. Finally, gyrification has been suggested to capture further morphometric subtleties of asymmetries [29, 58] but has not yet been assessed as a functional-behavioural structural asymmetry substrate.

Given available supporting evidence, our hypotheses and analyses emphasise the PT in terms of leftward surface-area asymmetry. In the supplementary material we also report exploratory analyses conducted on PT thickness and gyrification, as well as HG and IFGop surface-area, thickness, and gyrification (the IFGtr is also included for clarification of dissociation from the IFGop; that is, to support its exclusion as a structural asymmetry substrate of speech-related processes and challenge the use of the classical Broca’s region definition).

#### Use of probabilistic tractography

Diffusion MRI, particularly diffusion tensor imaging (DTI) was a revolution to the field of cognitive neuroscience [74]. Considerable critique for its capacity to reliably and validly measure white matter has since arisen, with a core issue being that the majority of voxels in the MRI sequence contain two or more fibre populations, often with different orientations, thereby distorting the output inferred fibre trajectory reconstruction [for review of this and other issues see 75]. Probabilistic tractography addresses uncertainty in fibre orientation(s) and using seed-based classification can provide a map of confidence values of connection across the brain [for more in-depth explanation see 76]. Streamlines reaching from voxels of a seed ROI that reach a target ROI can be divided by the total voxel count of the seed ROI to give an indirect indication of their connecting fibre count (although sometimes referred to as “connection strength” we use the more conservative term “[streamline] connectivity”). This approach has been previously validated and shown to predict behavioural task performance and functional laterality [77, 78]. When the variable of interest is asymmetry of physical size of a pathway [79], it is possible to use the output streamline pathway for volumetric analysis. Although not without its own limitations, probabilistic tractography shows promising consistency with “gold standard” invasive procedures limited to post-mortem and animal studies [80].

#### Use of individualized region-of-interest masks for tractography

Marked interindividual variability exists for our ROI [see 81, 82]. A major shortcoming of many previous dMRI studies as such has been using generic ROI templates in determining white matter connectivity. An advantage of the present paradigm is the use of FreeSurfer surface-based analysis to extract individualized ROI masks for tractography in FSL 5.0.1 (www.fmrib.ox.ac.uk/fsl).

#### Use of planum temporale transcallosal pathway

The corpus callosum and PT have both received considerable attention as substrates to functional lateralisation of speech [8, 22]. However, their relationship has been relatively unexplored, particularly with use of contemporary structural MRI techniques.

Post-mortem findings indicated that increased leftward surface area-asymmetry of the PT is related to reduced callosal connections between the left and right PT [83]. Thus, this study indicated a negative correlation of PT surface area asymmetry and transcallosal connectivity. To our awareness, the only dMRI study (using DTI measures) addressing the link of transcallosal connectivity to speech-related behavioural task performance indicated a positive relationship with greater radial diffusivity [more sensory/inhibitory connections; 84]. Thus, this study indicated a positive correlation of speech-related processing ability and transcallosal connectivity. These two studies then ostensibly present a contradiction towards our overarching hypothesis that PT surface area asymmetry and speech-related processing ability positively correlate (and consequently uniformly correlate to transcallosal connectivity).

To resolve the contradiction, we can consider that transcallosal fibres have microstructural and functional variances, with measures of the former being sensitive to method of measurement. Decreased radial diffusivity [measured by Elmer et al.; 84] has been proposed to reflect myelin thickness rather than axon number [inverse to axial diffusivity; 85, 86]. These larger-diameter fibres function more towards sensory/inhibitory signalling, whilst chance and colleagues’ method of measuring callosal size has been suggested to reflect density of small-diameter axons, which function more towards associative/excitatory signalling [87–89]. Therefore, the findings may represent a dynamic excitatory-inhibitory (rather than singular) function of the corpus callosum via reduced small (associative/excitatory) fibres and increased large (sensory/inhibitory) fibres. Greater presence of transcallosal inhibitory/sensory fibres [cf. Elmer et al.; 84] and lesser excitatory/associative fibres fits with the division-of-labour hypothesis of greater asymmetry and performance ability. However, this interpretation is speculative and requires validation, particularly using dMRI methods less susceptible to confounds such as crossing fibres inherent in DTI. Probabilistic tractography is such a measure, and generally sensitive to small diameter (associative/excitatory) fibres [76, 90].

#### Use of planum temporale to inferior frontal gyrus pars opercularis portion of arcuate fasciculus

Structural definition of the AF has some controversy in the existing literature [91–94]. A consolidating perspective identifies two temporal-frontal portions [95], the second of which (the “direct” pathway) captures the “classical” AF [cf. Geshwind; 96]. There is substantial evidence of leftward structural asymmetry of the AF, particularly for this direct frontal-temporal portion [e.g., 21, 97-99]. Such asymmetry has been implicated as a substrate of leftward language lateralisation in terms of cortical activation and behavioural performance in healthy adult and child populations [100–103]. Indeed, this portion of the AF was considered the major “speech pathway” in the classical model [50].

Whilst the inferior premotor cortex has more been proposed as a predominate site of tract termination for temporal-frontal pathways of the AF [104], this aligns with the posited “indirect” pathway of Friederici [95]. The IFGop (though not IFGtr) is identified as another robust point of termination [71, 79], and aligns with the “direct” temporal-frontal pathway. Starting points intending to capture Wernicke’s area have been as liberal as the entire inferior parietal and lateral temporal cortices [78]. However, they are generally more restricted.

Indeed, the PT alone has been equated with Wernicke’s area elsewhere [105]. Again, the primary focus for this study in broad is the relationship of structural asymmetries in speech-related processes and their lateralisation. Thus, we selected the PT and IFGop as the target ROI given supporting evidence for both in terms of leftward morphometrical asymmetry [9, 29], prior use as ROI in dMRI measurement of the AF [56, 71, 106] and a functional role in speech-related processing [107, 108].

#### Use of age, sex, and total brain volume as asymmetry control variables

The role of age and sex as confounds for structural asymmetries is mixed [as is the case with functional lateralities; 18]. Both factors have been shown to produce differences at the whole brain level for cortical volume [109]. [9] found that increasing age in adults (with at least 20 years difference of age range) produced reduced asymmetry at the whole hemispheric level yet increased leftward asymmetry in perisylvian regions. The authors also reported males to show more surface-area asymmetry than females at the regional level. Sex differences of cortical thickness in auditory-related cortical regions have also been reported [110]. Overall brain size has been proposed as a more general underlying factor for both measures, producing non-proportional differences in cortical surface-area and gyrification [60]. Indeed, [9] reported that intracranial volume showed mixed correlations depending on the regions and measure of interest. Given the potential influence of age, sex, and brain size, we included these measures as control variables.

#### Hypotheses

The primary aims of this study were to provide supporting evidence for relationships of morphometry and connectivity in speech-related cortex with behavioural measures of functional lateralisation of speech and abilities in speech-related acoustic processing. Specifically, we sought to support the hypothesis that the PT would be indicated as a structural substrate of language lateralisation (as measured by NF-DLT REA) and RTP for non-speech stimuli (as measured by GDT). Identified substrates were leftward surface area asymmetry, interhemispheric (PT-to-PT) transcallosal connectivity, and asymmetry of intrahemispheric connectivity with the IFGop (i.e., AF tracts from “Wernicke” to “Broca” regions). We also aimed to provide evidence for consideration and implementation of developing methodology in assessing morphometry and connectivity. To keep appropriate focus to our primary set of hypotheses however, these analyses and the related discussion are presented as supplementary material.

To our knowledge, RTP ability as measured by GDT task performance has not been previously directly assessed with PT surface-area or thickness. Greater leftward PT surface-area has been linked to better RTP ability [and AST-related leftward functional lateralisation; 67] and leftward PT thickness asymmetry has been related to better longer temporal processing [and AST-related rightward functional lateralisation; 69] using other tasks. We hypothesised that increased leftward PT surface-area asymmetry would correlate with increased RTP ability (i.e., lower GDT). Leftward volume asymmetry of the PT has some evidence of relating to leftward language lateralisation as reflected by REA in the behavioural DLT [111, 112]. We hypothesised that surface-area would more strongly demonstrate this relationship and thus that leftward asymmetry would correlate with REA. Assuming that VOT sensitivity would underlie the REA [46], we also predicted that controlling for RTP would nullify this structural asymmetry-functional laterality relationship.

Regarding interhemispheric connectivity, we firstly hypothesised that connectivity as indicated by PT-to-PT streamline connections would inversely correlate with leftward PT surface-area asymmetry. This was built on the premise that a previously reported inverse correlation from post-mortem analysis reflected a reduction of associative myelinated axons (excitatory small diameter, which our tractography measure is sensitive towards), thereby providing a structural architecture for division-of-labour processing [83, 113]. We thus also hypothesised that PT-to-PT transcallosal connectivity would negatively correlate to leftward language lateralisation (as measured by NF-DLT REA) and positively with RTP (as inversely measured by GDT). Furthermore, we predicted that the hypothesised association of leftward PT surface-area asymmetry to GDT and DLT would be at least partially reduced by controlling for transcallosal connectivity [cf. Chance; 83, 113].

Finally, regarding intrahemispheric connectivity, we hypothesised that the PT to IFGop portion of the AF as outlined by probabilistic tractography would show leftward volume asymmetry [22, 106]. The degree of this asymmetry was also predicted to positively relate to PT surface-area asymmetry, leftward language lateralisation and RTP [8, 114]. Again, partial overlap of the grey matter and white matter asymmetry effects with behavioural measures was hypothesised. In particular, controlling for surface-area asymmetry of the PT and IFGop was expected to at least partially reduce associations of AF volume asymmetry with GDT and NF-DLT REA.

## Methods

### Participants

Sixty-three eligible right-handed participants (40 female) aged 18 to 46 years (mean 24 years, median 22 years) volunteered for the study. Fifty participants were undergraduate psychology students who obtained course credit for their involvement, whilst the remaining 13 were members of the general public reimbursed AUD$60 for incurred incidental costs and time. Recruitment and data collection occurred from July 1^st^ 2015 until October 1^st^ 2016. Procedures were approved by the University of Newcastle Human Research Ethics Committee (H-276-0806) and all participants provided informed written consent. Although 71 individuals volunteered, eight were excluded due to raised hearing thresholds in one or both ears based on audiometric testing (>20 dB SPL between 500 and 6000 Hz), having a first-degree relative with schizophrenia, non-right-handedness [assessed using the Edinburgh Handedness Inventory; 115], reported use of recreational drugs or excessive alcohol consumption based on the Australian National Health and Medical Research Council guidelines, or unusable data in the GDT task. Two participants did not undergo the second session, leaving 61 usable MRI data-sets.

### Procedure

The measures and data reported in this study were collected alongside others not reported here (see appendix 8). Following the provision of written informed consent, participants were allocated to a project code that specified the order of task completion (left-ear conditions first or right-ear conditions) and whether the sounds they heard would be comprised of a low frequency leading marker and high frequency trailing marker or vice versa. Order for dichotic listening was also specified. This ensured that ear and marker frequency were counter-balanced across participants. For this phase, all participants attended one three-hour appointment during which they were required to complete (in order) a hearing test, the Edinburgh Handedness Inventory [115] to measure degree of right-handedness, a brief questionnaire to assess musical experience and monolingualism, the dichotic listening task, a gap-detection task to determine gap detection thresholds and the Wechsler Test of Adult Reading [116]. In the second session of two hours, participants completed the MRI scanning, reaction time tasks, and listening span task. The GDT and DLT measures relevant to this chapter are described below.

#### Gap-detection thresholds

Between-channel gap thresholds were estimated according to the procedure of Todd and colleagues [49], using a two-interval two-alternative forced choice procedure. A “gap” and a “no-gap” stimulus were presented in random order on each of 100 trials. Participants were instructed to click the left mouse button to indicate the first sound, or right click to indicate the second contained a gap. The initial gap size was fixed at 100 ms and was then varied according to performance using a 3-down-1-up psychophysical rule converging on an estimate on the smallest gap that could be detected on 79.4% of trials [117]. The change in gap size across trials was increased or reduced as a factor (1.2) of the preceding gap size (in 0.25 ms increments). That is, after three correct responses the gap size would decrease by a factor of 1.2 × the previous gap size and would increase by 1.2 × the previous gap size when an error was made. All stimuli were delivered monaurally and the task was repeated two times per participant [as per Lister et al., who averaged over two estimates obtained after a practice block; 118]. A third run was administered when attention or motivation confounds were evident with a discrepancy larger than 10ms between the two blocks. The most consistent two scores were then averaged to obtain estimate left and right-ear thresholds. Visual feedback was given following each trial and the threshold for each attempt was calculated as the geometric mean over the final eight reversal points.

The sounds for GDT task were presented over Sennheiser 280 PRO headphones. They were constructed digitally using MAT-LAB at a sampling frequency of 22.05 kHz. At the first stage a 132300-point Fast Fourier Transform (FFT) buffer with a sampling rate of 44,100 Hz was created (FFT resolution = 0.33 Hz). Each FFT buffer comprised 1201 individual FFT components (real and imaginary complex pairs). An inverse FFT specifying the frequency composition of the sound was then applied to each buffer to create each sound signal. Two sets of sounds were created, one with high frequency band leading markers (2200–2600 Hz) and low frequency band trailing markers (3400–3800 Hz) and one with low frequency band leading markers and high frequency band trailing markers. Ten versions of each sound were created with different randomization seeds and the gap-detection tasks sampled randomly from these ten each time a sound of a given gap size was required by the task. All sounds were presented at 75dB SPL calibrated using a Brüel & Kjær modular precision sound analyser (Type 2260) and sound level calibrator (Type 4231).

#### Dichotic listening task

The DLT procedure was conducted as outlined by Hugdahl [119, 120]. Three stimulus sets were used, each comprising 36 CVS pairs, consisting of six CVSs (/ba/, /da/, /ka/, /ta/, /pa/, /ga/). The pairs were spoken dichotically by a male voice at approximately 70dB SPL over Sennheiser 280 PRO headphones. The non-forced (NF) condition used the first stimulus set. Participants were shown a printout of the six CVSs and instructed that they should verbally respond to each dichotic presentation by reporting aloud the CVS they “heard best or most clearly [without thinking about the response] directly after it has been presented”. Thirty pairs were dichotomous (e.g., /ka/ to the left-ear, /ba/ to the right-ear), and six pairs were diotic (e.g., /ka/ to the left and right-ear). Diotic presentations, which should be easily reported, were to control for validity of participant responses (e.g., difficulties with stimulus perception, attention or motivation, technical issues etc.). The forced-right (FR) and forced-left (FL) conditions used the second and third stimulus sets in counterbalance. Order of completing these two conditions was also counterbalanced, but always followed NF. In FR/L, participants were instructed that they would repeat the same task as previously in NF. However, they were required to “only listen to and report the sound in the right/left-ear … ignore [sound in] the other ear”. Each condition took approximately four minutes to complete, including instructions (12 minutes total).

#### Structural Imaging

Structural imaging occurred in the second session, either before or after completion of a reaction time task and listening span task. The imaging protocol used a 3T Siemens Magnetom Prisma scanner and involved five acquisitions relevant to the presented data: (1) scout scan (14 sec) for scanner positioning, (2; 3) two T1 MPRAGE scans (∼5mins each) averaged for morphometrical analyses and inclusion/exclusion parameters for probabilistic tractography, (4) an inverse anterior-to-posterior diffusion weighted scan (∼14mins; for artifact reduction) [121], and (5) posterior-to-anterior diffusion weighted scan (∼7 mins). A T2 Flair (∼6 mins), WIP900 MP2RAGE (∼9 mins), WIP9000 memprage (∼6 mins) were also collected at the end of the scanning procedure for analyses not reported in the present thesis, taking total scanning time to approximately 52 mins. A brief safety screening questionnaire and induction was performed by the radiographer prior to scanning (∼8 mins). All scans were reviewed by a radiologist for any clinical abnormality that may compromise data; none such were observed. The relevant acquisitions and their processing are elaborated below.

#### Morphometrical imaging acquisition and preprocessing

Two high-resolution whole brain T1 anatomical images (MPRAGE; TR = 2000 ms, TE = 3.5 ms, Flip Angle = 7°, voxel size = 1mm isotropic, GRAPPA acceleration factor 2) were acquired. These anatomical images were averaged, segmented, and then inflated to the cortical surface using FreeSurfer [122]. Once inflated, the cortical surfaces of both hemispheres were co-registered to a spherical coordinate system [123]. The PT, HG, IFGop, and IFGtr were identified and parcellated automatically using the Destrieux et al. neuroanatomical atlas [53]. Volume, surface-area, and thickness of each ROI were extrapolated from the segmentation for statistical analysis [124, 125]. Local gyrification indexes were also extrapolated in FreeSurfer 6.0 by computing the ratio of outer surface-area to overall surface-area of each region [126]. An asymmetry index was calculated for each measurement of each ROI for every participant.

#### Diffusion-weighted imaging acquisition and preprocessing

A 32-channel array head coil with maximum gradient strength of 80 mT/m was used for the diffusion-weighted data acquisition. To correct for direction biases both an anterior-to-posterior (A-P) and inverse posterior-to-anterior (P-A) phase encoding sequence was acquired. Each used spin-echo planar imaging (TR = 6 sec, TE = 61 ms, 64 axial slices, resolution 2 x 2 x 2 mm, GRAPPA acceleration factor = 2). Diffusion weighting was isotropically distributed along 64 directions (bvalue = 1000 s/mm^2^, A-P excitations = 2; P-A excitation = 1). Note that high angular resolution of the diffusion weighting directions yields robust estimation of the fibre directions by increasing the signal-to-noise ratio and reducing directional bias. Three data sets with no diffusion weighting (b0) were acquired in A-P (start, middle, and finish), whilst two b0 images (start and finish) were acquired in P-A. These images served as an anatomical reference for offline motion correction.

Using FSL 5.0.1 (www.fmrib.ox.ac.uk/fsl) all baseline b0 images were aligned to a reference b0 image to estimate motion correction parameters using rigid-body transformations implemented in FLIRT [127]. The resulting linear transformation matrices were combined with a global registration to the T1 anatomy computed with the same method. The gradient direction for each volume was corrected using the rotation parameters. The transformation matrices were applied to the diffusion weighted images, and the corresponding acquisitions and gradient directions were averaged to improve signal-to-noise ratio [128].

#### Tractography

Diffusion image preprocessing and analyses were done using FSL 5.0.1. In accordance with Behrens et al. [129], estimation of tracts was conducted using probabilistic tractography. A probabilistic fibre tracking approach was chosen, using 10,000 tract-following samples at each voxel with a curvature threshold of 0.2. A dual-fibre model as implemented in bedpostX (FSL 5.0.1) was used. Dual-fibre models account for crossing fibres [128] therefore yielding more reliable results compared to single-fibre models. All tractography was done in each participant’s native space (un-normalized) data, and resulting maps were warped into standard space (using the MNI 1 mm isotropic brain as reference) for cross-participant averaging and comparison.

Estimation of connectivity between ROI was conducted using individualised seed masks for each participant. That is, the surface-based PT and IFGop ROI used in the mMRI analyses were converted to volume files using FreeSurfer. This was done by converting the Destrieux and colleagues [53] neuroanatomical atlas annotations produced in the FreeSurfer auto-segmentation to label files using the white matter surface as boundaries. These were then converted to volume files using a 0.5 fill threshold, binarised, and trimmed to a ribbon mask to prevent spill-over. The left and right ventricles and overall left and right hemispheres were likewise converted into volume files to be used respectively as exclusion and inclusion masks. Visual inspection ensured that all masks aligned with the brain volume and diffusion files in FSL.

For PT-to-PT connectivity, seed-based classification was done by first thresholding the images such that only voxels with at least ten samples are kept [130, 131]. Next, voxel values were converted into proportions to provide streamline connectivity indexes (i.e., “white matter tract strength”) such that the value at each voxel becomes the number of samples reaching the target mask for that image, divided by the number of samples from the seed mask. Seed-based classification was done from the left to right PT. Since directionality cannot be inferred from tractography analyses, seed-based classification was also done from the right to left PT and both directions were averaged. Left and right hemisphere white matter volumes were used as inclusion masks (to ensure fibres were interhemispheric), and masks of the ventricles were used as exclusion masks (to ensure fibres crossed via the corpus callosum).

For the AF, tractography was conducted using the same methods as above but using individualised PT masks as seeds and IFGop as targets. The total streamline pathways connecting the seeds and targets were converted into volume files, the values of which were then extracted. Again, to control for directionality tractography was also run from IFGop-to-PT and both directions were averaged.

#### Analyses

All statistical analyses were conducted using *SPSS Statistics 25.0* [132].

## Results

### Gap detection thresholds

The mean difference between the two GDT estimates averaged to generate the threshold measure for each individual was M = 3.12 ms, SD = 2.82 ms for left-ear and M = 3.8 ms, SD = 2.85 ms for the right-ear. A paired sample t-test showed that there was no significant difference between the average GDT obtained for the left (M = 17.87 ms, SD = 7.06 ms) and the right-ear (M = 17.82 ms, SD = 6.77 ms). A total threshold was created by averaging measurement of both ears. The distribution of averaged GDT scores is demonstrated in Figure 1. The overall mean was M = 17.85 ms, SD = 6.52, with a close median of 16.62 ms, and range of 36.14 ms (highest = 44.87 ms, lowest = 8.73).

**Figure 1.**
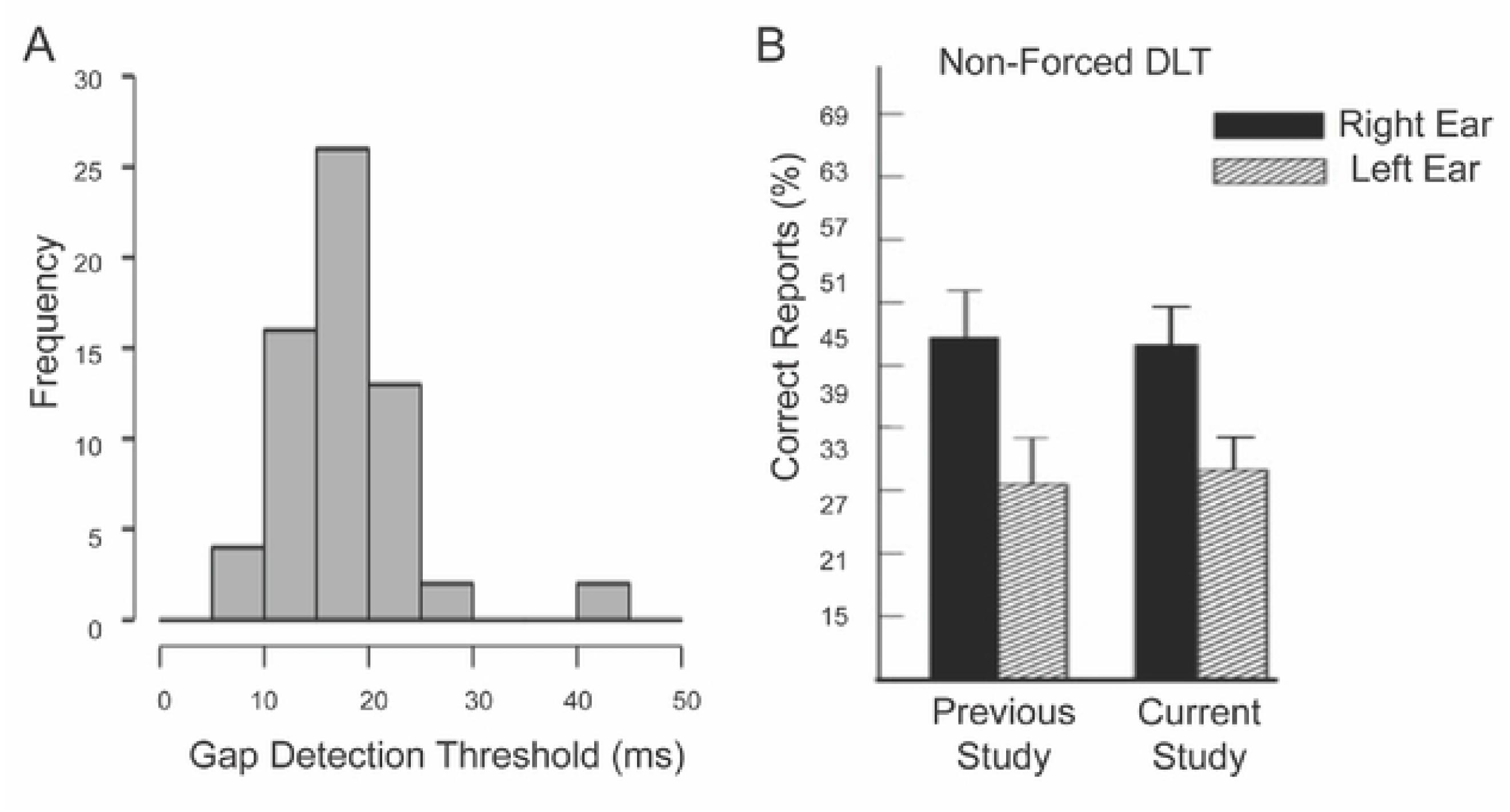
A. Frequency distribution of gap detection thresholds. B Dichotic listening task results of current study compared to previous by Hugdahl et al, 2009 for the non-forced attention condition. Note. Figure reproduced and amended with permission. Error bars = SD/2.

**Figure 2:**
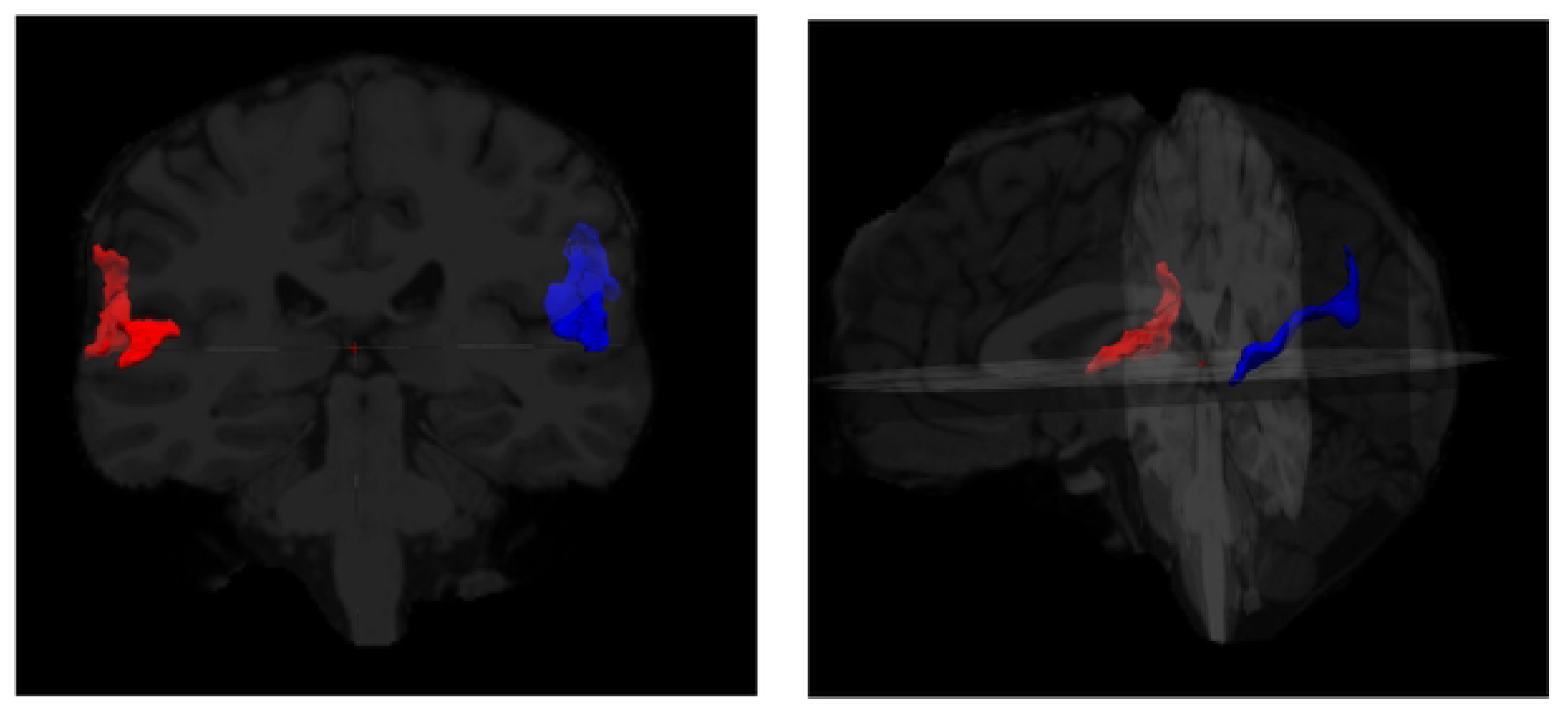
Example of individualised PT masks (generated in FreeSurfer). Note. Left PT white matter boundary in blue, right PT white matter boundary in red.

The gap detection thresholds of our study were remarkably similar to Todd et al. [49] with the latter average (∼19 ms) and standard deviation (∼7 ms) showing little discrepancy from the present study result. This is encouraging given that the task procedure was an exact replication (although participant demographics of age and sex ratio differed). It should be noted that two outliers (>2 standard deviations from mean) produced non-normality of the distribution, however parametric tests were implemented due to a precedent in the literature supporting the robustness (and advantages) of these measures with reasonable samples sizes [133, 134]. These two high threshold outliers were included, due to evidence of being representative of the population distribution [49]. Consistent with hypotheses, an independent samples t-test showed no significant differences of GDT between males and females, and bivariate correlations revealed no significant relationship between age and GDT. Lack of sex effects affirms the findings of Todd et al. [49]. Although the authors found an effect of age, our sample intentionally excluded the aged population (65 years and over) where these effects are expected to emerge. Thus, age and sex were not considered primary influencing factors of GDT in our study.

### Dichotic listening task

Dichotic listening task performance was explored by viewing the percentage of correct responses to right-ear and left-ear stimulus presentations, as well as a language laterality index derived from their difference (i.e., 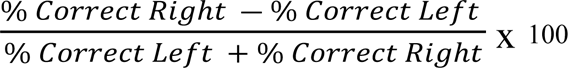; positive value = leftward lateralised). Pearson correlations (two-tailed) were performed to assess for relationships between age and DLT performance for each ear. As expected, no significant correlations or notable trends emerged. An independent samples t-test also showed no effects of sex on DLT performance.

### Non-forced condition

To assess language lateralisation using the non-forced DLT, paired sample t-tests were conducted to explore differences of ear across the sample. As hypothesised, a significant REA emerged (see figure 1 with a greater percentage of correct responses for right-ear (M = 46.98%, SD = 7.98) than left-ear (M = 35.56%, SD = 7.31) presentations, t(62) = 8.06, p < .001, and a mean REA of 13.85 (SD = 14.57).

Within the sample, fifty-four (∼86%) participants in our sample showed a REA. A paired sample t-test demonstrated that this was a significantly greater number of correct right-ear responses (M = 48.83, SD = 6.38), than left-ear responses (M = 34.57, SD = 7), t(53) = 11.08, p <.001. Eight participants exhibited a LEA. One participant showed no ear advantage. A significantly higher overall response accuracy (i.e., mean of left- and right-ear) was shown for those with a REA (M = 41.7, SD = 4.75), than those with a left-ear advantage (M = 37.7, SD = 6.84), t(60) = 2.09, p = .041. As shown in figure 1, NF-DLT outcomes of our sample were similar to those of Hugdahl and colleagues [31; n = 60; age range = 20-30 years].

### Rapid temporal processing and language lateralisation

Bivariate correlations (one-tailed) were conducted between participant GDT scores and DLT-NF to test our hypothesis that superior RTP (i.e., lower GDT) would be related to greater dispositional language laterality (i.e., NF-REA). Contradictory to our predictions, no relationship was evident between GDT and laterality in NF-DLT.

### Morphometrical measures

The surface area of the left and right PT for all participants were extracted using individualised masks (generated in Freesurfer; see figure #) to create an asymmetry index of surface area (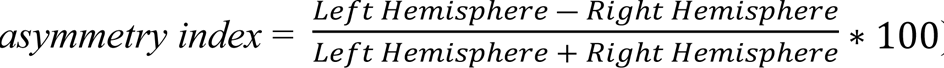). A paired samples t-test indicated that leftward asymmetry was statistically significant across the group (left PT: M = 667.54mm^2^, SD = 109.13 mm^2^, right PT: M = 596.1mm^2^, SD = 100.06 mm^2^), *t*(60) = 4.39, *p* <.001. Our sample showed a normal distribution for asymmetry index centred on the right (as expected; with positive values indicating leftward asymmetry), with a mean index of 5.62 (SD = 9.94). Total brain volume was also extracted, showing a mean size of 1,5572,285.82 mm^3^ (SD = 186,937.93 mm^3^; min = 1,068,925 mm^3^, max = 1,968,870 mm^3^). An independent samples t-test supported that males showed larger total brain volume than females, *t*(59) = 6.53, *p* <.001.

### Structural Asymmetries and Behavioural Abilities

#### Rapid temporal processing and planum temporale

Partial bivariate correlations (one-tailed; controlling for sex, age, and total brain volume) were conducted to assess for predicted relationships between measures of leftward PT surface-area asymmetry with better RTP ability as measured by GDTs. As predicted, a weak negative correlation was found for PT surface-area asymmetry, *r* = −.28, *p* = .017.

Thus, greater leftward asymmetry of PT surface-area indeed seemed related to behavioural rapid temporal acuity as measured with non-speech stimuli and in the direction of better ability. Our findings extend previous surface-based analyses showing improved temporal-cues-based CVS processing with greater left PT surface-area size [67].

#### Language lateralisation and planum temporale

Partial bivariate correlations (one-tailed; controlling for sex, age, and total brain volume) were conducted to assess for predicted relationships between leftward PT surface-area asymmetry with stronger leftward language lateralisation as measured by REA in the NF-DLT. As predicted, a weak positive correlation was found for PT surface-area asymmetry, *r* = .245, *p* = .032 (larger surface area asymmetry, better NF-DLT). Thus, greater leftward asymmetry of PT surface-area indeed seemed related to behavioural leftward language lateralisation.

### Probabilistic tractography of the planum temporale

#### Planum temporale transcallosal connections

As illustrated in figure 3, our use of probabilistic tractography constrained by PT seed and target masks showed an anatomically viable streamline output. Elmer, Hänggi [84] have previously shown a similar streamline path using the same paradigm and software for probabilistic tractography. Importantly however, rather than use generic masks as with the prior study (two spherical templates positioned in identical yet flipped space), our own tractography used individualised seed and target masks of each participant’s own left and right PT parcellation. Again, PT-to-PT connections were quantified using seed-based streamline connectivity. volume) were conducted to explore the expected relationships of PT-to-PT connectivity with our measures of PT surface-area asymmetry, language lateralisation, and RTP ability. As predicted, our results showed an inverse correlation of PT-to-PT connectivity and leftward PT surface-area asymmetry, *r* = −.309, *p* = .01. Further in line with the hypotheses, a significant inverse relationship emerged with leftward language lateralisation as measured by the NF-DLT REA, *r* = −.365, *p* = .003. In contrast to our hypotheses, behavioural RTP ability as measured by GDTs did not show a meaningful or statistical relationship with connectivity strength (see figure 4).

**Figure 3:**
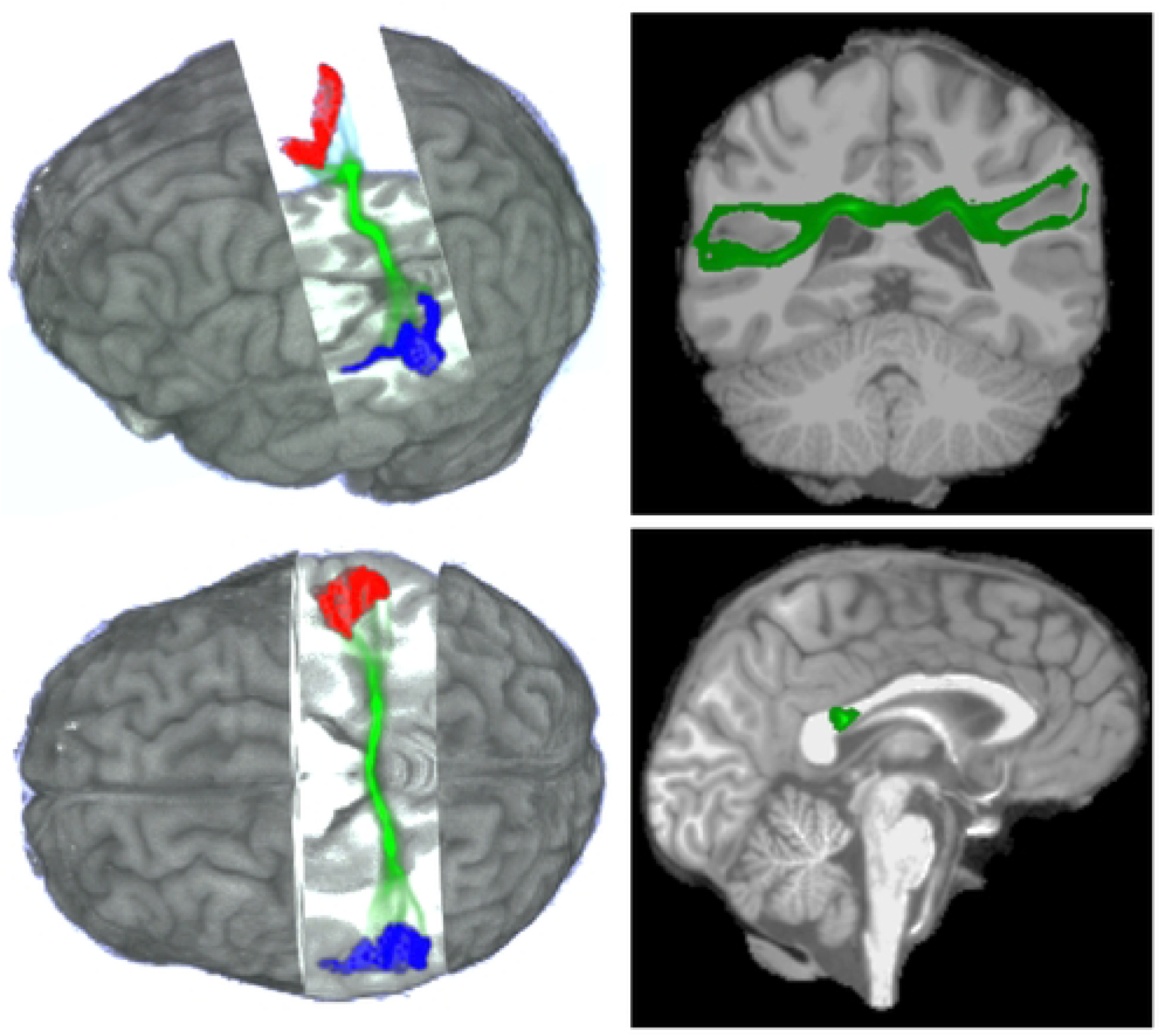
Example of tractography streamline pathways (generated with FSL) using individualised PT masks (generated in FreeSurfer). Note. Left PT white matter boundary in blue, right PT white matter boundary in red, transcallosal pathway in green.

**Figure 4:**
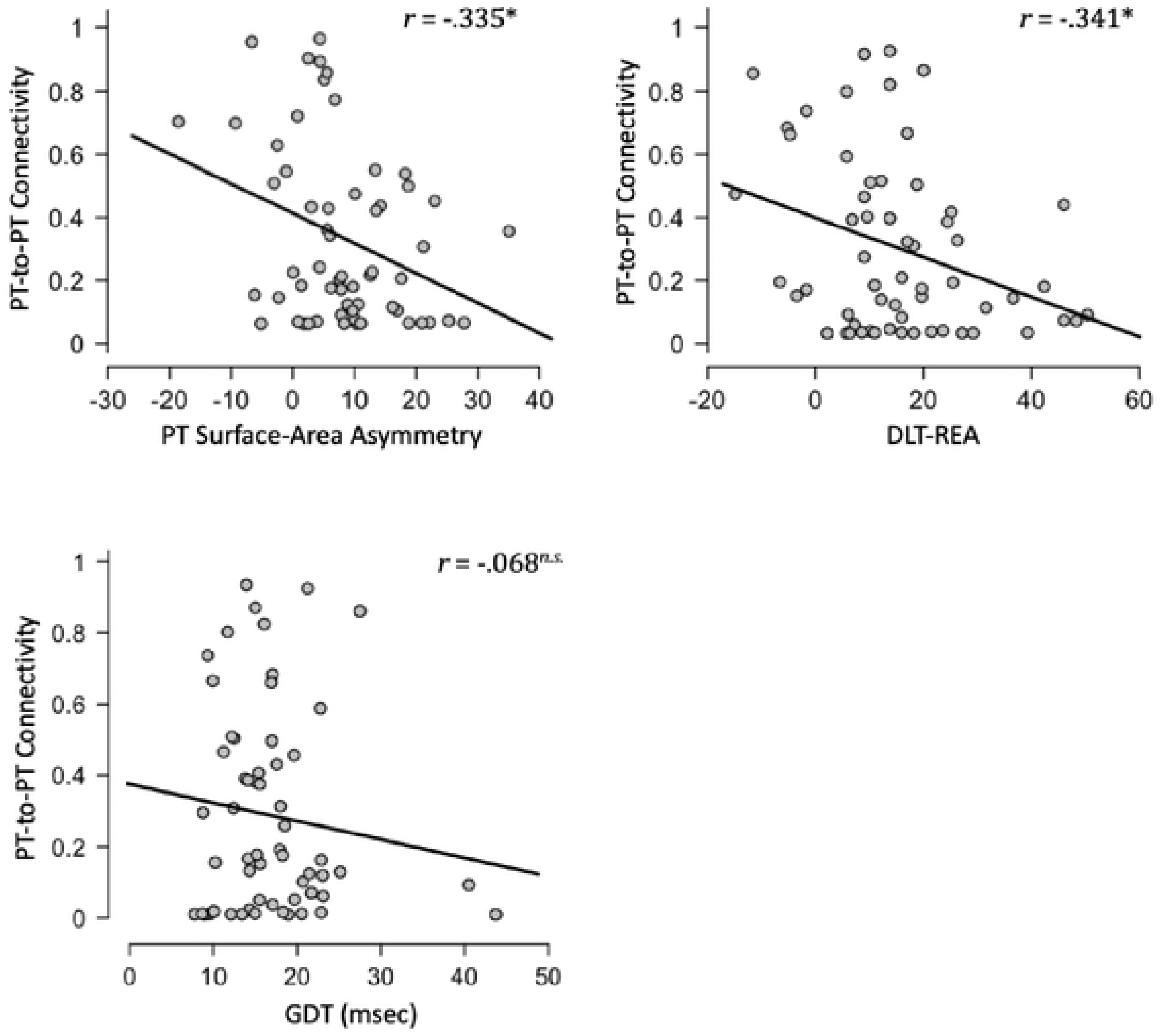
Scatterplots of PT-to-PT connectivity with leftward surface-area asymmetry of PT, right-ear advantage of dichotic listening performance (DLT-REA; i.e., leftward language lateralisation), and gap detection threshold (GDT; i.e., rapid temporal processing ability).

#### Arcuate fasciculus asymmetry

As illustrated in figure 5, our use of probabilistic tractography constrained by PT-to-IFGop seed and target masks showed an anatomically viable streamline output. This replicated several previous studies where probabilistic tractography has been applied for assessing the AF [135]. Our tractography followed previous use of the PT and IFGop as starting and termination points intended to capture contemporary analogues of Wernicke’s and Broca’s areas [56] and maintain our core focus on regions of asymmetry. We used volume of the tractography streamline pathways as the quantitative measure.

**Figure 5:**
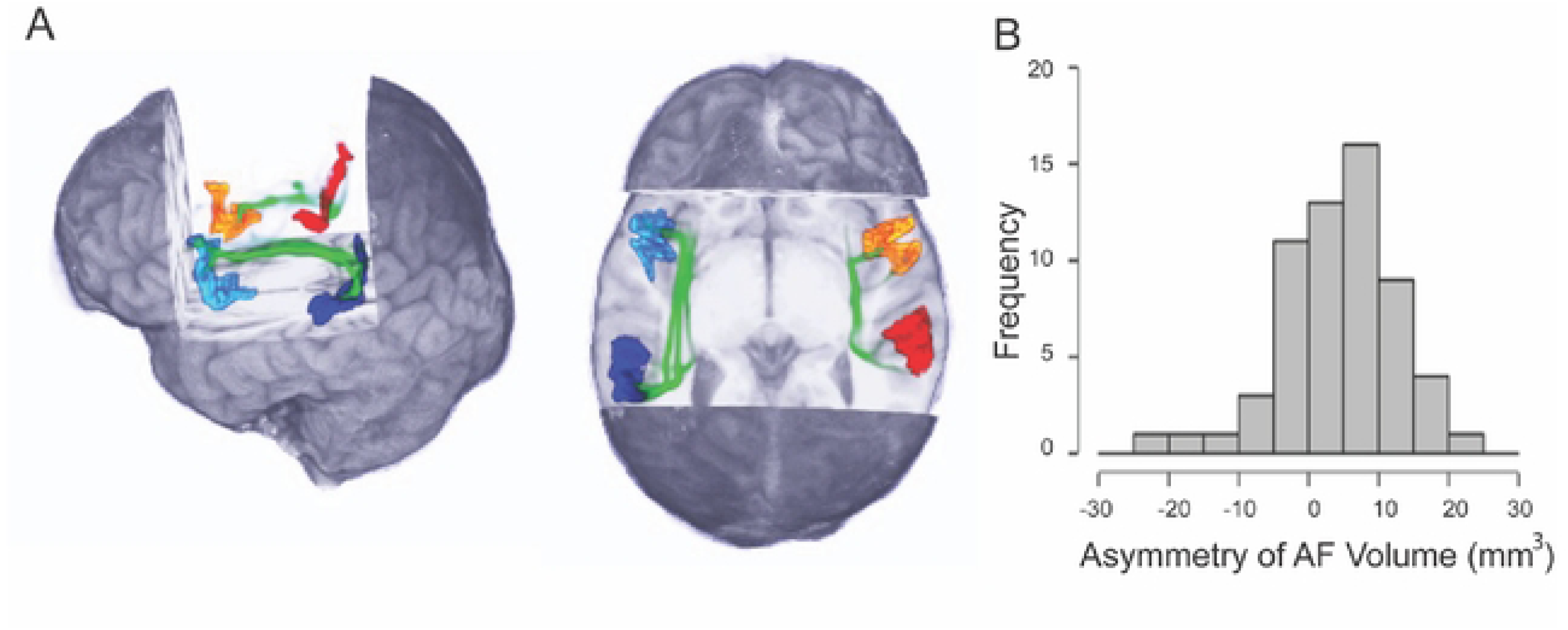
A. Example of tractography streamline pathways (generated with FSL) of the direct temporal-frontal AF using individualised PT and IFGop masks (generated in FreeSurfer). Note. White matter boundary of left PT in dark blue, left IFGop in cyan, right PT in red, right IFGop in orange, AF pathways in green. B. Frequency distribution of direct temporal-frontal arcuate fasciculus (AF) asymmetry indexes (positive value = leftward asymmetry).

Paired samples t-tests were conducted to address our hypothesis for a significant leftward AF volume asymmetry. As predicted, the left AF (M = 61299 mm^3^; SD = 1206 mm^3^) was significantly larger in volume than the right AF (M = 56596 mm^3^; SD = 1256 mm^3^) across participants, *t*(59) = 3.626, *p* < .001. As per our morphometrical data, an asymmetry index was calculated from the left and right AF volume values (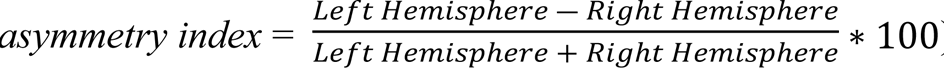). As shown in figure 5B, our sample showed a mostly normal distribution, centering on leftward asymmetry.

Partial bivariate correlations (one-tailed; controlling for sex, age, and total brain volume) were conducted to explore the expected relationships of direct temporal-frontal AF volume asymmetry with behavioural measures. In contrast to our predictions, no relationship was evident for AF volume asymmetry with either RTP ability (measured via GDTs) or language laterality (measured via NF-DLT REA).

#### Inter-relations of structural measures with behavioural measures

We predicted that various structural measures would overlap in their relationship to RTP and language lateralisation. Further partial bivariate correlations (one-tailed) were therefore conducted to assess overlap of our morphometrical, connective, and behavioural measures. As shown in table 4, indeed many structural measures related to each other as well as behavioural measures. These analyses controlled for sex, age, and total brain volume, but also alternatively PT-to-PT connectivity, direct temporal-frontal AF volume asymmetry, PT surface-area asymmetry, DLT REA, and GDT.

**Table 4.** Structural and Behavioural Measure Relationshiop.

|  |  | GDT | NF-DLT<br>REA | PT<br>AI | PT-to-PT<br>Connectivity | AF<br>AI |
| --- | --- | --- | --- | --- | --- | --- |
| GDT | Pearson's $r$ | — | | | | |
| | $p$-value | — | | | | |
| NF-DLT<br>REA | Pearson's $r$ | 0.060 | — | | | |
| | $p$-value | 0.321 | — | | | |
| PT AI | Pearson's $r$ | **-0.208** | **0.258 \*** | — | | |
| | $p$-value | **0.054** | **0.022** | — | | |
| PT-to-PT<br>Connectivity | Pearson's $r$ | -0.120 | **-0.329 \*\*** | **-0.328 \*\*** | — | |
| | $p$-value | 0.183 | **0.005** | **0.006** | — | |
| AF AI | Pearson's $r$ | -0.109 | 0.080 | **0.447 \*\*\*** | **-0.321 \*\*** | — |
| | $p$-value | 0.204 | 0.271 | **< .001** | **0.007** | — |

Note . \* $p < .05$ , \*\* $p < .01$ , \*\*\* $p < .001$ , one-tailed. GDT = gap detection threshold (inverse rapid temporal processing ability), NF-DLT REA = Non-forced Dichotic Listening Task Right Ear Advantage (leftward language lateralisation), PT = planum temporale, IFGop = inferior frontal gyrus pars opercularis, AF = arcuate fasciculus, AI = asymmetry index.

As predicted, NF-DLT REA showed relationship to both PT-to-PT connectivity (negative correlation) and PT surface-area asymmetry (positive correlation). The relationships of PT-to-PT connectivity survived corrections: NF-DLT REA (controlling for PT surface-area asymmetry), *r* = −.321, *p* = .01; PT surface-area asymmetry (controlling for NF-DLT REA), *r* = −.370, *p* = .003. The relationship of PT surface-area asymmetry and NF-DLT REA did not survive controlling for PT-to-PT connectivity, *r* = −.089, *p* = .265. We had predicted that the hypothesised relationship of language lateralisation and PT asymmetry would be at least partially attributable to PT-to-PT connectivity [cf. Chance; 113]. Our findings showed a completely eliminated relationship however. Controlling for PT-to-PT connectivity for the previously reported positive association of leftward PT surface-area asymmetry and RTP ability (as measured inversely by between-channel GDT) only strengthened the relationship, *r* = −304, *p* = .012. Thus, the present findings suggest that although leftward surface-area asymmetry of the PT and reduced PT-to-PT connectivity relate to leftward language lateralisation (and each other), the degree of connectivity appears to be the predominant structural factor. However, only PT surface-area asymmetry was a related structural factor for behavioural RTP.

## Discussion

### Dichotic listening and gap detection measures

Our outcomes for NF-DLT and GDT showed close resemblance to those of Hugdahl et al. [31] and Todd et al. [49], the respective prior studies on which our behavioural measures of REA and RTP were based. This is encouraging for interpretation of our outcomes of relationships of these measures to our structural measures.

### Planum temporale

Leftward asymmetry of the PT is one of the most robustly reported structural asymmetries for the human brain [136]. Our findings again evince this phenomenon and add to the growing evidence that such asymmetry is best attributed to cortical surface-area [28, 29, 57–60], which has been explained as a consequence of greater microcolumnar number, width, and spacing [83, 113, 137].Thus, leftward surface-area asymmetry of the PT indeed appears to be a general phenomenon in the human brain (see supplementary material for evidence that this constituent measure may also best represent prior reports of a volumetric asymmetry).

Our hypothesis that leftward surface-area asymmetry would positively relate to RTP behavioural ability (i.e., lower GDT) was affirmed. There is ample research supporting the PT as an anatomical substrate of functional lateralisations and behavioural ability in speech related processes [66, 107, 138–140]. Indeed, the columnar width and spacing differences underlying surface-area differences [83, 141] are a purported substrate for leftward PT lateralisation for finer-resolution “narrowing” processing and rightward lateralisation for broader “widening” processing [142–144], which has been proposed to underlie biases for speech and music, respectively [14, 145, 146]. However, studies directly linking structural asymmetry with functional lateralisation and performance are limited. Indeed, to our knowledge no other study has reported RTP as measured by GDTs with any measure of PT asymmetry. Left PT surface-area has been related to processing of rapid temporal information in a CVS and reduced-spectrum CVS-analogue categorisation task [67]. Ours is the first study to directly show a relationship of leftward PT surface-area asymmetry with acoustic RTP ability.

Our hypothesis that leftward surface-area asymmetry would positively relate to language lateralisation as measured by a free recall DLT was also supported. That is, those who showed a stronger REA also showed stronger leftward asymmetry. Prior studies relating DLT and PT asymmetry are mixed in findings and mostly limited to volumetric analyses. A relationship between leftward PT volume asymmetry and REA in DLT has been shown in a dyslexic sample, but not matching controls [111], whilst another study demonstrated the relationship in healthy controls, but only in right-handed males [112].

Greve, Van der Haegen [28] found that leftward language lateralisation measured using functional MRI response in a DLT was not significantly related to a leftward surface-area asymmetry of the PT. However, a region overlapping part of the PT with the lateral aspect of the superior temporal gyrus did show such an association. Our study did show a significant relationship of behavioural response as measured by REA with leftward surface-area asymmetry across the sample. Thus, although replication is required, our findings suggest that cortical surface-area is indeed more sensitive than volume to structural-behavioural relationships. Moore and colleagues [147] recently showed correlation of left PT activation (measured via fMRI) to stimuli in a NF-DLT for children without listening difficulties, yet a (non-significant trend) pattern for those with. Interestingly, only subtle differences for behavioural DLT performance were evident between the groups. Thus, in comparing our findings to those of Greve and colleagues, it is also apparent that behavioural and neurophysiological DLT approaches vary, especially in their sensitivity to structural asymmetry.

In contrast to our predictions, although a positive relationship of leftward PT surface-area asymmetry was found for both RTP ability and language lateralisation, controlling for the former did not nullify the relationship of the latter. That is, our hypothesis that RTP would underlie the asymmetry-REA relationship was not supported. Manipulation of rapid temporal cues in the VOT of CVS has marked effects on DLT performance [for in depth review see 46]. However, our DLT paradigm did not likely sufficiently emphasise VOT as the core cue for CVS recognition, thus possibly explaining the lack of relationship between the two behavioural measures and their relationships to PT surface-area asymmetry.

### Connections of the Planum Temporale

#### Planum temporale transcallosal connections

Our use of probabilistic tractography constrained by PT seed and target masks showed an anatomically viable transcallosal streamline output. This replicates the previous work of Elmer and colleages [84]. Importantly, however, our own tractography used individualised seed and target masks of each PT. The previous study used generic masks (two spherical templates positioned in identical yet flipped space), imposing inherent confounds when assessing structures that are (as is the point in asymmetry studies) non-homogenous in terms of morphology.

Our findings affirmed an inverse relationship of PT-to-PT transcallosal connectivity with leftward PT surface-area asymmetry and behavioural language lateralisation (as measured via NF-DLT REA). Indeed, as discussed above, surface-area asymmetry and language lateralisation positively related. However, controlling for connectivity eliminated this relationship. In a post-mortem study, Chance and colleagues [83] showed that the left PT was greater in surface-area than the right PT, attributable to both greater microcolumn count (specifically in men) and microcolumn width and spacing. Greater columns count was also observed in the HG, though width and spacing was not. Indeed, the authors proposed that columnar spacing was the relevant factor for transcallosal connectivity and lateralisation of speech processing. Our own results give support for this proposal, with leftward PT surface-area relating to both PT-to-PT connectivity and behavioural language laterality.

We were unable to clarify the contributions of columnar count and spacing to our observed leftward PT surface-area asymmetry and its relation to functional lateralisation of speech. However, a recent study by Ocklenburg and others [148] gives strong support to emphasis of columnar architecture. The authors reported that the cortical columns of the left PT indeed have more width radially and in terms of spacing, yet are also more arborised [see also 143, 149, 150]. In brief, the left PT shows a more intracolumnar-connective structure than the right PT, which shows a more intercolumnar-connective structure. This gives a cytoarchitectonic analogue to the lateralised functional intrahemispheric processing of the left hemisphere, and interhemispheric processing of the right hemisphere [151–153].

Ocklenburg et al. [148] also reported functional lateralisation using a passive NF-DLT auditory evoked potential paradigm (controlling for VOT differences of CVSs). The component of interest was the N1, which the authors described as the earliest lateralised auditory evoked potential component indexing orientation towards and perception of verbal stimuli. Specifically, N1 latency was used as an index of psychophysiological sensitivity. The authors inferred that grey matter microstructure (i.e., neurite/columnar factors) was primary for leftward lateralisation of N1 response to CVSs. This was based on a lack of relationship of N1 latency response to macrostructural grey matter (PT cortical thickness and volume) and white matter (AF volume). Surprisingly however, PT surface-area and PT-to-PT connectivity were not reported, despite the authors proposing their significance in another recent paper [8].

Qin and colleages [27] assessed PT macrostructural (surface area and thickness) and microstructural (myelin content, neurite density, and neurite orientation dispersion) properties of the PT with functional lateralisation (measured using functional MRI) in speech perception and comprehension tasks. The authors reported a significant positive relationship of PT surface area (but not thickness) with lateralisation of speech comprehension. Interestingly, they concluded that microstructural features are more important than macrostructural features for assessing structural correlates of functional lateralisation of speech, given that these measures related to activity for both speech perception and comprehension. Notably, such relationships of microstructural features seemed selective to morphological subtypes of the PT (based upon HG gyrification patterns and subsequent delineation), whereas the effect of surface area and speech comprehension activity was more robust (unaffected by subtype).

Taken together with our own findings, it is apparent that grey matter macrostructure (surface-area), microstructure (density and dispersion patterns of myelin, neurite, and microcolumns etc.), and connectivity can each related to DLT processing and speech processing more broadly. Microstructure and tract connectivity may indeed underlie macrostructure relationships, yet further research conjunctively yet directly assessing each is still valuable. Continuing developments of microstructural measurement methods are demonstrating increasing feasibility and informativeness of such conjunctive studies [154–156]

In contrast to predictions, our findings did not show a relationship of RTP ability with PT-to-PT connectivity. Both Ocklenburg et al. [148] and Chance [113] propose that the leftward microstructural asymmetry of the PT provides an architecture for leftward lateralised RTP and attribute this as a primary substrate of speech and language lateralisation. Neither provided a direct measure of rapid temporal acuity in their work however. Again, our study did show a positive relationship of RTP ability with leftward cortical surface-area asymmetry, which was only strengthened by controlling for PT-to-PT connectivity. Thus, whilst PT-to-PT connectivity was the primary structural factor for behavioural language lateralisation, leftward PT surface-area asymmetry was the primary structural factor for behavioural RTP ability.

#### Direct frontal-temporal arcuate fasciculus connections

Our study replicated several previous studies where probabilistic tractography has been applied for assessing the AF, often using the IFGop as the point of termination [135]. Seed masks have been as liberal as the entire inferior parietal and lateral temporal cortices [78], yet generally more restricted. Again, the PT and IFGop have been previously used as starting and termination points [56] and maintained our core focus on regions of asymmetry. Consistent with these studies was a clear leftward asymmetry, particularly AF volume as reported by Ocklenburg [22].

In contrast to our hypotheses, leftward volume asymmetry of the PT-to-IFGop (direct temporal-frontal) portion of the AF did not show any relationship to behavioural measures of RTP or language lateralisation. Ocklenburg [22] previously reported a relationship of DLT performance with left AF volume, though not asymmetry specifically. However, the authors have previously proposed that such intrahemispheric asymmetry is a core substrate of lateralisation of speech-related processes [8, 22]. This perspective shifted in the authors most recent publication, with the proposal that neurite connectivity, but not the AF, is a core structural substrate of leftward lateralisation of speech-related processes. Our study also challenges the notion of leftward volume asymmetry of the direct temporal-frontal AF as a core substrate of speech-lateralisation (specifically behavioural RTP and language lateralisation). Indeed, controlling for asymmetry of the seed and target ROI eliminated the asymmetry altogether. However, this is in context of only our specific means of measuring the AF. Starting/terminating and inclusion/exclusion masks of the AF, demarcation of its various portions (i.e., temporal-frontal, temporal parietal-frontal, direct, indirect etc.), and methods of its quantification (e.g., volume, fractional anisotropy, connectivity etc.) are widely variable [135].

#### Structural substrates of rapid temporal processing and language lateralisation

To our knowledge no other study has conjunctively assessed behavioural RTP ability, language lateralisation, PT morphometry asymmetries, PT-to-PT connectivity, and direct temporal-frontal AF volume asymmetry. The key findings of our own study are that leftward PT surface-area asymmetry appears to be the primary structural correlate of behavioural RTP, and PT-to-PT connectivity is the primary structural correlate of behavioural language lateralisation.

Elmer and others [84] used probabilistic tractography to identify the PT-to-PT transcallosal pathway and then conduct DTI-based statistics (i.e., fractional anisotropy, axial diffusivity, & radial diffusivity). The study also included behavioural speech-related categorisation tasks using CVS and non-speech CVS-analogue stimuli. The authors found only a relationship for radial diffusivity, which inversely correlated with performance for both speech and non-speech analogue identification. Lower radial diffusivity has been [contentiously; see 157] proposed to reflect higher myelin “integrity”. More accurately, it (indirectly) reflects degree or thickness of myelination and axon diameter [85]. Thinness and thickness of myelin sheathing is relative for adaptivity and therefore thinner myelin only reflects integrity issues when of pathological cause or consequence [158–162]. These caveats considered, the findings may suggest a relationship of speech and non-speech processing with greater (sensory/inhibitory) interhemispheric connections, which typically have thicker myelin sheathing [87–89]. Thus, performance on speech-related categorisation tasks may be facilitated by more PT-to-PT sensory and inhibitory connections. As proposed by Elmer and colleagues, this involves RTP ability. Though again our study showed no relationship of PT-to-PT connectivity with a between-channel GDT measure of RTP acuity, our findings do potentially suggest that less transcallosal associative connections facilitate increased language lateralisation in a behavioural DLT task. Indeed, corpus callosum size in general has been shown to inversely relate to NF-DLT REA [163]. Leftward PT surface-area asymmetry also related to behavioural language lateralisation in our study, while the correlation did not survive correction for PT-to-PT connectivity. Behavioural RTP correlated with leftward PT surface-area asymmetry only, including after controlling for PT-to-PT connectivity.

Synthesizing the findings and perspectives of the current study with those of Chance [83, 113], Ocklenberg [8, 148], and Elmer [67, 84] and their colleagues it is clear that macrostructural and microstructural grey and white matter factors of the PT relate to functional lateralisation of behavioural RTP and speech-processing. The specifics of how these structural substrates overlap is much less clear. In general, it is apparent that the greater microcolumnar spacing, width, and aborisation of the left PT may facilitate the stereotypical rapid temporal, linear, and template matching processing of the left hemisphere. As counterpart to this intracolumnar architecture, the intercolumnar architecture of the right PT may facilitate (optimally) slower, non-linear, and integrative processing, which is characteristic of the right-hemisphere in general [13, 14, 164, 165]. The division-of-labour afforded to each hemisphere in its specialised processing style may be mediated by the degree of associative and sensory/inhibitory transcallosal connections (positively and negatively respectively). The extent of relevance and involvement of these structural factors requires further investigation and likely holds the same caveat as their functional correlates: it depends on the context.

#### Limitations and future developments

Some limitations regarding methodology of the present study have been addressed in the above discussion. Our methods of inference of white matter connectivity and volume, as with all dMRI studies, are indirect. Although this is an issue for all diffusion studies, it is one that is underacknowledged [74, 75, 166]. Our use of seed-based classification with individualised masks for tracts previously identified using post-mortem and tractography analyses did optimise the validity of our tracts [80]. However, careful interpretation is required for the currently nascent methodological field. Indeed, consilience of methods is perhaps the best safe-guard. That is, combining multiple diffusion approaches can ensure confidence in the reliability and validity of results. The emerging technique of ‘g-ratio’ for example can offer a better index of ratio of axon to myelin sheath [87, 167]. This would considerably facilitate clarification of the relevant role of associative and sensory/inhibitory connections in PT-to-PT and direct temporal-frontal AF pathways. As well as g-ratio, other newer methods are in constant development and although still undergoing validation indicate promising improvements [154–156]. Our acquisition protocol did not facilitate such analysis approaches, an important consideration for future dMRI studies.

Our mMRI measures have better methodological validity, with a more solid extant literature for consistency of *in vivo* and post-mortem morphology measures [53, 64, 65, 168]. Indeed, a particular strength of the current study was the use of cortical surface-area in place of cortical volume [as well as thickness and local gyrification; see supplementary material; 29, 58]. Where this could be strengthened however, is with more specific parcellations of ROI. An emerging focus in the literature is on the morphological variability of both the HG [169, 170] and PT [82]. Functionally distinct regions of the PT have been previously identified using manual segmentation [138, 171]. Use of developing automatic surface-based sub-segmentation software [e.g., 172] may allow more specific analyses of the asymmetry of these PT sub-regions and their involvement in speech-related lateralisation. Furthermore, as emphasised by Qin and colleagues [27] microstructural asymmetries of the PT (intracortical myelin, content, neurite density, & neurite orientation dispersion) also show relationships with functional lateralisation for speech-processing. Although these authors claim microstructure factors to be superior variables of interest, their results also showed micro/macro to be differentially sensitive to morphological variances – a finding that we suggest is strongly meriting of conjunctive use and exploration of both microstructure and macrostructure.

Regarding our behavioural measurement of functional lateralisation, Westerhausen and Samuelsen [173] published an improved version of the DLT after our data collection. This update minimises the relevance of higher cognitive functions on task performance in order to obtain stimulus-driven laterality estimates, derived more directly from stop-consonant vowel (CV) syllables as the relevant cue. This does not invalidate our use of the prior measures (with over 50 years of use prior) but would be an improvement for future studies.

Finally, another important improvement would be to balance lateralisation and ability for linguistic and acoustic behavioural measures. We assessed the former pairs only with the DLT and GDT. A valuable extension would be the inclusion of a task measuring lateralisation of non-linguistic RTP and behavioural ability for linguistic detection (e.g., speech-in-noise task).

### Conclusion

This study continues understanding of how structural asymmetries of speech-related cortex are associated with behavioural measures of language-lateralisation and rapid temporal processing ability. Morphometrical MRI measures included the surface-area (as well as thickness and gyrification; see supplementary material) of left and right PT (as well as HG, IFGop, and IFGtr; see supplementary material). We affirmed recent literature that suggests cortical volume is better represented by surface-area. Asymmetry indexes of the PT showed leftward laterality to be “normal” (though variable) in our sample. The relationship of leftward PT surface-area asymmetry with behavioural measures was also reported, showing an expected positive association with both RTP ability (inversely measured using GDTs) and language lateralisation (measured using NF-DLT REA). Use of probabilistic tractography to determine PT-to-PT connectivity showed an inverse relationship to language lateralisation, which eliminated the relationship with surface-area asymmetry. This was interpreted to reflect PT-to-PT connectivity as the predominant structural factor for language lateralisation though improved division-of-labour via reduced transcallosal associative connections.

Drawing from prior research, the leftward PT surface-area asymmetry was posited to result from microcolumnar architecture that is facilitative of RTP. Overall, these findings support the existence of relationships of speech-relevant structural asymmetries, functional lateralisation, and behavioural ability, yet also demonstrate their complexity.

## References

1. Broca P. Remarques sur le si ege de la facult e du langage articul e, suivies d’une observation d’aph emie (perte de la parole). Bull Soc anatomique de Paris. 1861;6:330-56.

2. Wernicke C. Der aphasische Symptomencomplex: eine psychologische Studie auf anatomischer Basis: Cohen and Weigert; 1874.

3. Geschwind N, Levitsky W. Human brain: left-right asymmetries in temporal speech region. Science. 1968;161(3837):186-7.

4. Clunies-Ross KL. An Exploration of Hemispheric Asymmetries in Temporal Processing During Middle Childhood and Adulthood and Their Relationship with Language and Reading During Development. Perth, WA: University of Western Australia; 2019.

5. Parviainen T, Helenius P, Salmelin R. Children show hemispheric differences in the basic auditory response properties. Hum Brain Mapp. 2019;40(9):2699-710.

6. Westerhausen R. A primer on dichotic listening as a paradigm for the assessment of hemispheric asymmetry. Laterality. 2019;24(6):740-71.

7. Giroud J, Trebuchon A, Schon D, Marquis P, Liegeois-Chauvel C, Poeppel D, et al. Asymmetric sampling in human auditory cortex reveals spectral processing hierarchy. PLoS Biol. 2020;18(3):e3000207.

8. Ocklenburg S, Friedrich P, Gunturkun O, Genc E. Intrahemispheric white matter asymmetries: the missing link between brain structure and functional lateralization? Rev Neurosci. 2016;27(5):465-80.

9. Kong X-Z, Mathias S, Guadalupe T, Abé C, Agartz I, Akudjedu TN, et al. Mapping Cortical Brain Asymmetry in 17,141 Healthy Individuals Worldwide via the ENIGMA Consortium. bioRxiv. 2017.

10. Angenstein N, Brechmann A. Division of labor between left and right human auditory cortices during the processing of intensity and duration. NeuroImage. 2013;83:1-11.

11. Badzakova-Trajkov G, Corballis MC, Haberling IS. Complementarity or independence of hemispheric specializations? A brief review. Neuropsychologia. 2016;93(Pt B):386-93.

12. Keller M, Neuschwander P, Meyer M. When right becomes less right: Neural dedifferentiation during suprasegmental speech processing in the aging brain. NeuroImage. 2019;189:886-95.

13. Scott SK, McGettigan C. Do temporal processes underlie left hemisphere dominance in speech perception? Brain and language. 2013;127(1):36-45.

14. Zatorre RJ, Gandour JT. Neural specializations for speech and pitch: moving beyond the dichotomies. Philosophical transactions of the Royal Society of London Series B, Biological sciences. 2008;363(1493):1087-104.

15. Boemio A, Fromm S, Braun A, Poeppel D. Hierarchical and asymmetric temporal sensitivity in human auditory cortices. Nature neuroscience. 2005;8(3):389-95.

16. Poeppel D. The analysis of speech in different temporal integration windows: cerebral lateralization as ‘asymmetric sampling in time’. Speech Communication. 2003;41(1):245-55.

17. Poeppel D, Assaneo MF. Speech rhythms and their neural foundations. Nat Rev Neurosci. 2020;21(6):322-34.

18. Bourke JD, Todd J. Acoustics versus linguistics? Context is Part and Parcel to lateralized processing of the parts and parcels of speech. Laterality. 2021:1-41.

19. Toga AW, Thompson PM. Mapping brain asymmetry. Nat Rev Neurosci. 2003;4(1):37-48.

20. Jäncke L, Liem F, Merillat S. Are language skills related to structural features in Broca’s and Wernicke’s area? European Journal of Neuroscience. 2021;53(4):1124-35.

21. Vernooij MW, Smits M, Wielopolski PA, Houston GC, Krestin GP, van der Lugt A. Fiber density asymmetry of the arcuate fasciculus in relation to functional hemispheric language lateralization in both right- and left-handed healthy subjects: A combined fMRI and DTI study. NeuroImage. 2007;35(3):1064-76.

22. Ocklenburg S, Schlaffke L, Hugdahl K, Westerhausen R. From structure to function in the lateralized brain: how structural properties of the arcuate and uncinate fasciculus are associated with dichotic listening performance. Neurosci Lett. 2014;580:32-6.

23. Yazbek S, Hage S, Mallak I, Smayra T. Tractography of the Arcuate Fasciculus in Healthy Right-Handed and Left-Handed Multilingual Subjects and Its Relation to Language Lateralization on Functional MRI. 2021.

24. Galaburda AM, Corsiglia J, Rosen GD, Sherman GF. Planum temporale asymmetry, reappraisal since Geschwind and Levitsky. Neuropsychologia. 1987;25(6):853-68.

25. Foundas AL, Leonard CM, Gilmore R, Fennell E, Heilman KM. Planum temporale asymmetry and language dominance. Neuropsychologia. 1994;32(10):1225-31.

26. Isler B, Giroud N, Hirsiger S, Kleinjung T, Meyer M. Bilateral age-related atrophy in the planum temporale is associated with vowel discrimination difficulty in healthy older adults. Hear Res. 2021;406:108252.

27. Qin P, Bi Q, Guo Z, Yang L, Li H, Li P, et al. Microstructural asymmetries of the planum temporale predict functional lateralization of auditory-language processing. 2023.

28. Greve DN, Van der Haegen L, Cai Q, Stufflebeam S, Sabuncu MR, Fischl B, et al. A surface-based analysis of language lateralization and cortical asymmetry. Journal of cognitive neuroscience. 2013;25(9):1477-92.

29. Chiarello C, Vazquez D, Felton A, McDowell A. Structural asymmetry of the human cerebral cortex: Regional and between-subject variability of surface area, cortical thickness, and local gyrification. Neuropsychologia. 2016;93(Pt B):365-79.

30. Hugdahl K. Fifty years of dichotic listening research - still going and going and. Brain Cogn. 2011;76(2):211-3.

31. Hugdahl K, Westerhausen R, Alho K, Medvedev S, Laine M, Hamalainen H. Attention and cognitive control: unfolding the dichotic listening story. Scandinavian journal of psychology. 2009;50(1):11-22.

32. Bethmann A, Tempelmann C, De Bleser R, Scheich H, Brechmann A. Determining language laterality by fMRI and dichotic listening. Brain research. 2007;1133(1):145-57.

33. Binder JR, Swanson SJ, Hammeke TA, Morris GL, Mueller WM, Fischer M, et al. Determination of language dominance using functional MRI A comparison with the Wada test. Neurology. 1996;46(4):978-84.

34. Hugdahl K, Carlsson G, Uvebrant P, Lundervold AJ. Dichotic-listening performance and intracarotid injections of amobarbital in children and adolescents: preoperative and postoperative comparisons. Archives of neurology. 1997;54(12):1494-500.

35. Rutten GJ, Ramsey NF, van Rijen PC, van Veelen CW. Reproducibility of fMRI- determined language lateralization in individual subjects. Brain and language. 2002;80(3):421-37.

36. Heinrich A, Alain C, Schneider BA. Within-and between-channel gap detection in the human auditory cortex. Neuroreport. 2004;15(13):2051-6.

37. Salthouse TA. The processing-speed theory of adult age differences in cognition. Psychological review. 1996;103(3):403.

38. Phillips DP, Taylor T, Hall S, Carr M, Mossop J. Detection of silent intervals between noises activating different perceptual channels: some properties of “central” auditory gap detection. The Journal of the Acoustical Society of America. 1997;101(6):3694-705.

39. Steinschneider M, Volkov IO, Noh MD, Garell PC, Howard III MA. Temporal encoding of the voice onset time phonetic parameter by field potentials recorded directly from human auditory cortex. Journal of neurophysiology. 1999;82(5):2346-57.

40. Frye RE, Fisher JM, Coty A, Zarella M, Liederman J, Halgren E. Linear coding of voice onset time. Journal of cognitive neuroscience. 2007;19(9):1476-87.

41. Pichora-Fuller MK, Schneider BA, Benson NJ, Hamstra SJ, Storzer E. Effect of age on detection of gaps in speech and nonspeech markers varying in duration and spectral symmetry. The Journal of the Acoustical Society of America. 2006;119(2):1143.

42. Elangovan S, Stuart A. Natural boundaries in gap detection are related to categorical perception of stop consonants. Ear and hearing. 2008;29(5):761-74.

43. Ross B, Schneider B, Snyder JS, Alain C. Biological markers of auditory gap detection in young, middle-aged, and older adults. PloS one. 2010;5(4):e10101.

44. Heimrath K, Kuehne M, Heinze HJ, Zaehle T. Transcranial direct current stimulation (tDCS) traces the predominance of the left auditory cortex for processing of rapidly changing acoustic information. Neuroscience. 2014;261:68-73.

45. Sulakhe N, Elias LJ, Lejbak L. Hemispheric asymmetries for gap detection depend on noise type. Brain and Cognition. 2003;53(2):372-5.

46. Arciuli J. Manipulation of voice onset time during dichotic listening. Brain Cogn. 2011;76(2):233-8.

47. Rimol LM, Eichele T, Hugdahl K. The effect of voice-onset-time on dichotic listening with consonant-vowel syllables. Neuropsychologia. 2006;44(2):191-6.

48. Trebuchon-Da Fonseca A, Giraud K, Badier JM, Chauvel P, Liegeois-Chauvel C. Hemispheric lateralization of voice onset time (VOT) comparison between depth and scalp EEG recordings. NeuroImage. 2005;27(1):1-14.

49. Todd J, Finch B, Smith E, Budd TW, Schall U. Temporal processing ability is related to ear-asymmetry for detecting time cues in sound: a mismatch negativity (MMN) study. Neuropsychologia. 2011;49(1):69-82.

50. Tremblay P, Dick AS. Broca and Wernicke are dead, or moving past the classic model of language neurobiology. Brain and language. 2016;162:60-71.

51. Friedrich P, Anderson C, Schmitz J, Schluter C, Lor S, Stacho M, et al. Fundamental or forgotten? Is Pierre Paul Broca still relevant in modern neuroscience? Laterality. 2019;24(2):125-38.

52. Fedorenko E, Blank IA. Broca’s Area Is Not a Natural Kind. Trends Cogn Sci. 2020;24(4):270-84.

53. Destrieux C, Fischl B, Dale A, Halgren E. Automatic parcellation of human cortical gyri and sulci using standard anatomical nomenclature. NeuroImage. 2010;53(1):1-15.

54. Yaakub SN, Heckemann RA, Keller SS, McGinnity CJ, Weber B, Hammers A. On brain atlas choice and automatic segmentation methods: a comparison of MAPER & FreeSurfer using three atlas databases. Sci Rep. 2020;10(1):2837.

55. Lechanoine F, Jacquesson T, Beaujoin J, Serres B, Mohammadi M, Planty-Bonjour A, et al. WIKIBrainStem: An online atlas to manually segment the human brainstem at the mesoscopic scale from ultrahigh field MRI. NeuroImage. 2021;236:118080.

56. Upadhyay J, Hallock K, Ducros M, Kim DS, Ronen I. Diffusion tensor spectroscopy and imaging of the arcuate fasciculus. NeuroImage. 2008;39(1):1-9.

57. Panizzon MS, Fennema-Notestine C, Eyler LT, Jernigan TL, Prom-Wormley E, Neale M, et al. Distinct genetic influences on cortical surface area and cortical thickness. Cereb Cortex. 2009;19(11):2728-35.

58. Hogstrom LJ, Westlye LT, Walhovd KB, Fjell AM. The structure of the cerebral cortex across adult life: age-related patterns of surface area, thickness, and gyrification. Cereb Cortex. 2013;23(11):2521-30.

59. Koelkebeck K, Miyata J, Kubota M, Kohl W, Son S, Fukuyama H, et al. The contribution of cortical thickness and surface area to gray matter asymmetries in the healthy human brain. Hum Brain Mapp. 2014;35(12):6011-22.

60. Meyer M, Liem F, Hirsiger S, Jancke L, Hanggi J. Cortical surface area and cortical thickness demonstrate differential structural asymmetry in auditory-related areas of the human cortex. Cereb Cortex. 2014;24(10):2541-52.

61. Fischl B, Rajendran N, Busa E, Augustinack J, Hinds O, Yeo BT, et al. Cortical folding patterns and predicting cytoarchitecture. Cereb Cortex. 2008;18(8):1973-80.

62. Zilles K, Palomero-Gallagher N, Amunts K. Development of cortical folding during evolution and ontogeny. Trends Neurosci. 2013;36(5):275-84.

63. Fuhrmeister P, Myers EB. Structural neural correlates of individual differences in categorical perception. Brain and language. 2021;215:104919.

64. Fischl B. FreeSurfer. NeuroImage. 2012;62(2):774-81.

65. Desikan RS, Ségonne F, Fischl B, Quinn BT, Dickerson BC, Blacker D, et al. An automated labeling system for subdividing the human cerebral cortex on MRI scans into gyral based regions of interest. NeuroImage. 2006;31(3):968-80.

66. Meyer M, Elmer S, Jancke L. Musical expertise induces neuroplasticity of the planum temporale. Annals of the New York Academy of Sciences. 2012;1252:116-23.

67. Elmer S, Hänggi J, Meyer M, Jäncke L. Increased cortical surface area of the left planum temporale in musicians facilitates the categorization of phonetic and temporal speech sounds. Cortex. 2013;49(10):2812-21.

68. Neuschwander P, Hanggi J, Zekveld AA, Meyer M. Cortical thickness of left Heschl’s gyrus correlates with hearing acuity in adults - A surface-based morphometry study. Hear Res. 2019;384:107823.

69. Liem F, Hurschler MA, Jäncke L, Meyer M. On the planum temporale lateralization in suprasegmental speech perception: Evidence from a study investigating behavior, structure, and function. Human Brain Mapping. 2014;35(4):1779-89.

70. Warrier C, Wong P, Penhune V, Zatorre RJ, Parrish T, Abrams D, et al. Relating structure to function: Heschl’s gyrus and acoustic processing. The Journal of neuroscience : the official journal of the Society for Neuroscience. 2009;29(1):61-9.

71. Kaplan E, Naeser MA, Martin PI, Ho M, Wang Y, Baker E, et al. Horizontal portion of arcuate fasciculus fibers track to pars opercularis, not pars triangularis, in right and left hemispheres: a DTI study. NeuroImage. 2010;52(2):436-44.

72. Schremm A, Noven M, Horne M, Soderstrom P, van Westen D, Roll M. Cortical thickness of planum temporale and pars opercularis in native language tone processing. Brain and language. 2018;176:42-7.

73. Keller SS, Roberts N, García-Fiñana M, Mohammadi S, Ringelstein EB, Knecht S, et al. Can the language-dominant hemisphere be predicted by brain anatomy? Journal of cognitive neuroscience. 2011;23(8):2013-29.

74. Wandell BA. Clarifying Human White Matter. Annu Rev Neurosci. 2016;39:103-28.

75. Jones DK, Knosche TR, Turner R. White matter integrity, fiber count, and other fallacies: the do’s and don’ts of diffusion MRI. NeuroImage. 2013;73:239-54.

76. Behrens TEJ, Sotiropoulos SN, Jbabdi S. MR diffusion tractography. Diffusion MRI (Second Edition): Elsevier; 2014. p. 429-51.

77. Forstmann BU, Keuken MC, Jahfari S, Bazin PL, Neumann J, Schafer A, et al. Cortico-subthalamic white matter tract strength predicts interindividual efficacy in stopping a motor response. NeuroImage. 2012;60(1):370-5.

78. Takaya S, Kuperberg GR, Liu H, Greve DN, Makris N, Stufflebeam SM. Asymmetric projections of the arcuate fasciculus to the temporal cortex underlie lateralized language function in the human brain. Front Neuroanat. 2015;9:119.

79. Wu J, Lu J, Zhang H, Zhang J, Mao Y, Zhou L. Probabilistic map of language regions: challenge and implication. Brain : a journal of neurology. 2015;138(Pt 3):e337.

80. Donahue CJ, Sotiropoulos SN, Jbabdi S, Hernandez-Fernandez M, Behrens TE, Dyrby TB, et al. Using Diffusion Tractography to Predict Cortical Connection Strength and Distance: A Quantitative Comparison with Tracers in the Monkey. The Journal of neuroscience : the official journal of the Society for Neuroscience. 2016;36(25):6758-70.

81. Flinker A, Korzeniewska A, Shestyuk AY, Franaszczuk PJ, Dronkers NF, Knight RT, et al. Redefining the role of Broca’s area in speech. Proceedings of the National Academy of Sciences of the United States of America. 2015;112(9):2871-5.

82. Tzourio-Mazoyer N, Mazoyer B. Variations of planum temporale asymmetries with Heschl’s Gyri duplications and association with cognitive abilities: MRI investigation of 428 healthy volunteers. Brain Struct Funct. 2017.

83. Chance SA, Casanova MF, Switala AE, Crow TJ. Minicolumnar structure in Heschl’s gyrus and planum temporale: Asymmetries in relation to sex and callosal fiber number. Neuroscience. 2006;143(4):1041-50.

84. Elmer S, Hänggi J, Jäncke L. Interhemispheric transcallosal connectivity between the left and right planum temporale predicts musicianship, performance in temporal speech processing, and functional specialization. Brain Structure and Function. 2016;221(1):331-44.

85. Song S-K, Sun S-W, Ramsbottom MJ, Chang C, Russell J, Cross AH. Dysmyelination Revealed through MRI as Increased Radial (but Unchanged Axial) Diffusion of Water. NeuroImage. 2002;17(3):1429-36.

86. Beaulieu C, Does MD, Snyder RE, Allen PS. Changes in water diffusion due to Wallerian degeneration in peripheral nerve. Magnetic Resonance in Medicine. 1996;36(4):627-31.

87. Bjornholm L, Nikkinen J, Kiviniemi V, Nordstrom T, Niemela S, Drakesmith M, et al. Structural properties of the human corpus callosum: Multimodal assessment and sex differences. NeuroImage. 2017;152:108-18.

88. Aboitiz F, Scheibel AB, Fisher RS, Zaidel E. Individual differences in brain asymmetries and fiber composition in the human corpus callosum. Brain research. 1992;598(1):154-61.

89. Aboitiz F, Scheibel AB, Fisher RS, Zaidel E. Fiber composition of the human corpus callosum. Brain research. 1992;598(1):143-53.

90. Assaf Y, Alexander DC, Jones DK, Bizzi A, Behrens TE, Clark CA, et al. The CONNECT project: Combining macro- and micro-structure. NeuroImage. 2013;80:273-82.

91. Catani M, Jones DK, ffytche DH. Perisylvian language networks of the human brain. Ann Neurol. 2005;57(1):8-16.

92. Glasser MF, Rilling JK. DTI tractography of the human brain’s language pathways. Cereb Cortex. 2008;18(11):2471-82.

93. Petrides M, Pandya DN. Distinct parietal and temporal pathways to the homologues of Broca’s area in the monkey. PLoS Biol. 2009;7(8):e1000170.

94. Thiebaut de Schotten M, Ffytche DH, Bizzi A, Dell’Acqua F, Allin M, Walshe M, et al. Atlasing location, asymmetry and inter-subject variability of white matter tracts in the human brain with MR diffusion tractography. NeuroImage. 2011;54(1):49-59.

95. Friederici AD. White-matter pathways for speech and language processing. Handb Clin Neurol. 2015;129:177-86.

96. Geschwind N. Disconnexion syndromes in animals and man. Brain : a journal of neurology. 1965;88(3):585-.

97. Propper RE, O’Donnell LJ, Whalen S, Tie Y, Norton IH, Suarez RO, et al. A combined fMRI and DTI examination of functional language lateralization and arcuate fasciculus structure: Effects of degree versus direction of hand preference. Brain Cogn. 2010;73(2):85-92.

98. Nucifora PGP, Verma R, Melhem ER, Gur RE, Gur RC. Leftward asymmetry in relative fiber density of the arcuate fasciculus. NeuroReport. 2005;16(8):791-4.

99. Powell HWR, Parker GJM, Alexander DC, Symms MR, Boulby PA, Wheeler-Kingshott CAM, et al. Hemispheric asymmetries in language-related pathways: A combined functional MRI and tractography study. NeuroImage. 2006;32(1):388-99.

100. Sreedharan RM, Menon AC, James JS, Kesavadas C, Thomas SV. Arcuate fasciculus laterality by diffusion tensor imaging correlates with language laterality by functional MRI in preadolescent children. Neuroradiology. 2015;57(3):291-7.

101. Yeatman JD, Dougherty RF, Rykhlevskaia E, Sherbondy AJ, Deutsch GK, Wandell BA, et al. Anatomical Properties of the Arcuate Fasciculus Predict Phonological and Reading Skills in Children. Journal of cognitive neuroscience. 2011;23(11):3304-17.

102. Lebel C, Beaulieu C. Lateralization of the arcuate fasciculus from childhood to adulthood and its relation to cognitive abilities in children. Hum Brain Mapp. 2009;30(11):3563-73.

103. Allendorfer JB, Hernando KA, Hossain S, Nenert R, Holland SK, Szaflarski JP. Arcuate fasciculus asymmetry has a hand in language function but not handedness. Hum Brain Mapp. 2016;37(9):3297-309.

104. Brown EC, Jeong JW, Muzik O, Rothermel R, Matsuzaki N, Juhasz C, et al. Evaluating the arcuate fasciculus with combined diffusion-weighted MRI tractography and electrocorticography. Hum Brain Mapp. 2014;35(5):2333-47.

105. Stigler KA, McDougle CJ. Chapter 3.1 - Structural and Functional MRI Studies of Autism Spectrum Disorders A2 - Buxbaum, Joseph D. In: Hof PR, editor. The Neuroscience of Autism Spectrum Disorders. San Diego: Academic Press; 2013. p. 251-66.

106. Matsumoto R, Okada T, Mikuni N, Mitsueda-Ono T, Taki J, Sawamoto N, et al. Hemispheric asymmetry of the arcuate fasciculus. Journal of Neurology. 2008;255(11):1703.

107. Zheng ZZ. The functional specialization of the planum temporale. J Neurophysiol. 2009;102(6):3079-81.

108. Hagoort P. Nodes and networks in the neural architecture for language: Broca’s region and beyond. Current opinion in neurobiology. 2014;28:136-41.

109. Takahashi R, Ishii K, Kakigi T, Yokoyama K. Gender and age differences in normal adult human brain: voxel-based morphometric study. Hum Brain Mapp. 2011;32(7):1050-8.

110. Im K, Lee JM, Lee J, Shin YW, Kim IY, Kwon JS, et al. Gender difference analysis of cortical thickness in healthy young adults with surface-based methods. NeuroImage. 2006;31(1):31-8.

111. Hugdahl K, Heiervang E, Ersland L, Lundervold A, Steinmetz H, Smievoll AI. Significant relation between MR measures of planum temporale area and dichotic processing of syllables in dyslexic children. Neuropsychologia. 2003;41(6):666-75.

112. Dos Santos Sequeira S, Woerner W, Walter C, Kreuder F, Lueken U, Westerhausen R, et al. Handedness, dichotic-listening ear advantage, and gender effects on planum temporale asymmetry--a volumetric investigation using structural magnetic resonance imaging. Neuropsychologia. 2006;44(4):622-36.

113. Chance SA. The cortical microstructural basis of lateralized cognition: a review. Frontiers in psychology. 2014;5:820.

114. Della Penna S, Brancucci A, Babiloni C, Franciotti R, Pizzella V, Rossi D, et al. Lateralization of dichotic speech stimuli is based on specific auditory pathway interactions: neuromagnetic evidence. Cereb Cortex. 2007;17(10):2303-11.

115. Oldfield RC. The assessment and analysis of handedness: the Edinburgh inventory. Neuropsychologia. 1971;9(1):97-113.

116. Hugdahl K. Dichotic Listening with CV Syllables: Manual. 1994.

117. Hugdahl K. Dichotic listening in the study of auditory laterality. In: Hugdahl K, Davidson RJ, editors. The Asymmetrical Brain. Cambridge, MA: MIT Press; 2003. p. 441– 76.

118. Schaer M, Cuadra MB, Schmansky N, Fischl B, Thiran JP, Eliez S. How to measure cortical folding from MR images: a step-by-step tutorial to compute local gyrification index. J Vis Exp. 2012(59):e3417.

119. IBM. SPSS Statistics for Mac. 25.0 ed. Armonk, NY: IBM Corp; 2017.

120. Skovlund E, Fenstad GU. Should we always choose a nonparametric test when comparing two apparently nonnormal distributions? Journal of Clinical Epidemiology. 2001;54(1):86-92.

121. Fagerland MW. t-tests, non-parametric tests, and large studies—a paradox of statistical practice? BMC Medical Research Methodology. 2012;12(1):78.

122. Catani M, Budisavljević S. Contribution of Diffusion Tractography to the Anatomy of Language. In: Johansen-Berg H, Behrens TE, editors. Diffusion MRI (Second Edition). San Diego: Academic Press; 2014. p. 511-29.

123. Shapleske J, Rossell S, Woodruff P, David A. The planum temporale: a systematic, quantitative review of its structural, functional and clinical significance. Brain Research Reviews. 1999;29(1):26-49.

124. Chance SA, Clover L, Cousijn H, Currah L, Pettingill R, Esiri MM. Microanatomical correlates of cognitive ability and decline: normal ageing, MCI, and Alzheimer’s disease. Cereb Cortex. 2011;21(8):1870-8.

125. Tremblay P, Deschamps I, Gracco VL. Regional heterogeneity in the processing and the production of speech in the human planum temporale. Cortex. 2013;49(1):143-57.

126. Shrem T, Deouell LY. Frequency-dependent auditory space representation in the human planum temporale. Front Hum Neurosci. 2014;8:524.

127. Tzourio-Mazoyer N, Crivello F, Mazoyer B. Is the planum temporale surface area a marker of hemispheric or regional language lateralization? Brain Struct Funct. 2017.

128. Galuske RAW, Schlote W, Bratzke H, Singer W. Interhemispheric Asymmetries of the Modular Structure in Human Temporal Cortex. Science. 2000;289(5486):1946.

129. Rodel M, Cook ND, Regard M, Landis T. Hemispheric dissociation in judging semantic relations: Complementarity for close and distant associates. Brain and language. 1992;43(3):448-59.

130. Harasty J, Seldon HL, Chan P, Halliday G, Harding A. The left human speech-processing cortex is thinner but longer than the right. Laterality. 2003;8(3):247-60.

131. Jung-Beeman M. Bilateral brain processes for comprehending natural language. Trends Cogn Sci. 2005;9(11):512-8.

132. Zatorre RJ, Belin P. Spectral and temporal processing in human auditory cortex. Cerebral Cortex. 2001;11(10):946-53.

133. Zatorre RJ, Belin P, Penhune VB. Structure and function of auditory cortex: music and speech. Trends in cognitive sciences. 2002;6(1):37-46.

134. Moore DR, Hugdahl K, Stewart HJ, Vannest J, Perdew AJ, Sloat NT, et al. Listening Difficulties in Children: Behavior and Brain Activation Produced by Dichotic Listening of CV Syllables. Frontiers in psychology. 2020;11:675.

135. Ocklenburg S, Friedrich P, Fraenz C, Schlüter C, Beste C, Güntürkün O, et al. Neurite architecture of the planum temporale predicts neurophysiological processing of auditory speech. Science Advances. 2018;4(7).

136. Seldon HL. Structure of human auditory cortex. I. Cytoarchitectonics and dendritic distributions. Brain research. 1981;229(2):277-94.

137. Seldon HL. Structure of human auditory cortex. II. Axon distributions and morphological correlates of speech perception. Brain research. 1981;229(2):295-310.

138. Iturria-Medina Y, Perez Fernandez A, Morris DM, Canales-Rodriguez EJ, Haroon HA, Garcia Penton L, et al. Brain hemispheric structural efficiency and interconnectivity rightward asymmetry in human and nonhuman primates. Cereb Cortex. 2011;21(1):56-67.

139. Gotts SJ, Jo HJ, Wallace GL, Saad ZS, Cox RW, Martin A. Two distinct forms of functional lateralization in the human brain. Proceedings of the National Academy of Sciences of the United States of America. 2013;110(36):E3435-44.

140. Caeyenberghs K, Leemans A. Hemispheric lateralization of topological organization in structural brain networks. Hum Brain Mapp. 2014;35(9):4944-57.

141. Palombo M, Ianus A, Guerreri M, Nunes D, Alexander DC, Shemesh N, et al. SANDI: A compartment-based model for non-invasive apparent soma and neurite imaging by diffusion MRI. NeuroImage. 2020;215:116835.

142. Mohammadi S, Callaghan MF. Towards in vivo g-ratio mapping using MRI: Unifying myelin and diffusion imaging. Journal of Neuroscience Methods. 2021;348.

143. Tian Q, Bilgic B, Fan Q, Ngamsombat C, Zaretskaya N, Fultz NE, et al. Improving in vivo human cerebral cortical surface reconstruction using data-driven super-resolution. Cereb Cortex. 2021;31(1):463-82.

144. Wheeler-Kingshott CA, Cercignani M. About “axial” and “radial” diffusivities. Magn Reson Med. 2009;61(5):1255-60.

145. Fields RD. Myelin—More than Insulation. Science. 2014;344(6181):264-6.

146. Glasser MF, Goyal MS, Preuss TM, Raichle ME, Van Essen DC. Trends and properties of human cerebral cortex: correlations with cortical myelin content. NeuroImage. 2014;93 Pt 2:165-75.

147. Bercury KK, Macklin WB. Dynamics and mechanisms of CNS myelination. Dev Cell. 2015;32(4):447-58.

148. Fields RD, Bukalo O. Myelin makes memories. Nature neuroscience. 2020;23(4):469-70.

149. Knowles J, Batra A, Xu H, Monje M. Adaptive and maladaptive myelination in health and disease. Nature reviews Neurology. 2022.

150. Westerhausen R, Hugdahl K. The corpus callosum in dichotic listening studies of hemispheric asymmetry: a review of clinical and experimental evidence. Neuroscience and biobehavioral reviews. 2008;32(5):1044-54.

151. McGilchrist I. Reciprocal organization of the cerebral hemispheres. Dialogues in Clinical Neuroscience. 2010;12(4):503-15.

152. Poeppel D. The neuroanatomic and neurophysiological infrastructure for speech and language. Current opinion in neurobiology. 2014;28:142-9.

153. Sotiropoulos SN, Zalesky A. Building connectomes using diffusion MRI: why, how and but. NMR Biomed. 2017.

154. Mancini M, Giulietti G, Dowell N, Spano B, Harrison N, Bozzali M, et al. Introducing axonal myelination in connectomics: A preliminary analysis of g-ratio distribution in healthy subjects. NeuroImage. 2017.

155. Glasser MF, Coalson TS, Robinson EC, Hacker CD, Harwell J, Yacoub E, et al. A multi-modal parcellation of human cerebral cortex. Nature. 2016.

156. Marie D, Maingault S, Crivello F, Mazoyer B, Tzourio-Mazoyer N. Surface-Based Morphometry of Cortical Thickness and Surface Area Associated with Heschl’s Gyri Duplications in 430 Healthy Volunteers. Front Hum Neurosci. 2016;10:69.

157. Tzourio-Mazoyer N, Maingault S, Panzieri J, Pepe A, Crivello F, Mazoyer B. Intracortical Myelination of Heschl’s Gyrus and the Planum Temporale Varies With Heschl’s Duplication Pattern and Rhyming Performance: An Investigation of 440 Healthy Volunteers. Cerebral Cortex. 2018.

158. Isenberg AL, Vaden KI, Jr., Saberi K, Muftuler LT, Hickok G. Functionally distinct regions for spatial processing and sensory motor integration in the planum temporale. Hum Brain Mapp. 2012;33(10):2453-63.

159. Dalboni da Rocha JL, Santoro R, Van de Ville D, Golestani N. New method for automatic suface-based segmentation of Heschl’s gyrus. Organization for Human Brain Mapping; Vancouver, BC2017.

160. Westerhausen R, Samuelsen F. An optimal dichotic-listening paradigm for the assessment of hemispheric dominance for speech processing. PloS one. 2020;15(6):e0234665.

161. Junes FV, Barragan E, Alvarez D, Dies P, Tobon SH. Wernicke’s area and broca’s area in functional connectivity of language. 2019.

162. Nasios G, Dardiotis E, Messinis L. From Broca and Wernicke to the Neuromodulation Era: Insights of Brain Language Networks for Neurorehabilitation. Behav Neurol. 2019;2019:9894571.

163. Sprung-Much T, Eichert N, Nolan E, Petrides M. Broca’s area and the search for anatomical asymmetry: commentary and perspectives. Brain Struct Funct. 2022;227(2):441-9.

164. Benjamini Y, Hochberg Y. Controlling the false discovery rate: a practical and powerful approach to multiple testing. Journal of the Royal statistical society: series B (Methodological). 1995;57(1):289-300.

165. Pearlson G, E Barta P, E Powers R, R Menon R, S Richards S, H Aylward E, et al. Ziskind-Somerfeld Research Award 1996. Medial and superior temporal gyral volumes and cerebral asymmetry in schizophrenia versus bipolar disorder1997. 1-14 p.

166. Lin Y, Zhang K, Li S, Li S, Jin J, Jin F, et al. Relationship Between Perisylvian Essential Language Sites and Arcuate Fasciculus in the Left Hemisphere of Healthy Adults. Neurosci Bull. 2017;33(6):616-26.

167. Xiang HD, Fonteijn HM, Norris DG, Hagoort P. Topographical functional connectivity pattern in the perisylvian language networks. Cereb Cortex. 2010;20(3):549-60.

168. Steinschneider M, Nourski KV, Fishman YI. Representation of speech in human auditory cortex: is it special? Hear Res. 2013;305:57-73.

169. Alho J, Green BM, May PJC, Sams M, Tiitinen H, Rauschecker JP, et al. Early-latency categorical speech sound representations in the left inferior frontal gyrus. NeuroImage. 2016;129:214-23.

170. Kang X, Herron TJ, Ettlinger M, Woods DL. Hemispheric asymmetries in cortical and subcortical anatomy. Laterality. 2015:1-27.

171. Gunturkun O, Ocklenburg S. Ontogenesis of Lateralization. Neuron. 2017;94(2):249-63.

172. Maingault S, Tzourio-Mazoyer N, Mazoyer B, Crivello F. Regional correlations between cortical thickness and surface area asymmetries: A surface-based morphometry study of 250 adults. Neuropsychologia. 2016;93(Pt B):350-64.

173. Griffiths TD, Warren JD. The planum temporale as a computational hub. Trends in Neurosciences. 2002;25(7):348-53.

174. Steinschneider M, Volkov IO, Fishman YI, Oya H, Arezzo JC, Howard MA, 3rd. Intracortical responses in human and monkey primary auditory cortex support a temporal processing mechanism for encoding of the voice onset time phonetic parameter. Cereb Cortex. 2005;15(2):170-86.

175. Steinschneider M, Nourski KV, Rhone AE, Kawasaki H, Oya H, Howard MA, 3rd. Differential activation of human core, non-core and auditory-related cortex during speech categorization tasks as revealed by intracranial recordings. Frontiers in neuroscience. 2014;8:240.

176. Keller SS, Highley JR, Garcia-Finana M, Sluming V, Rezaie R, Roberts N. Sulcal variability, stereological measurement and asymmetry of Broca’s area on MR images. J Anat. 2007;211(4):534-55.

## Supplementary Material

The primary aims of the present study were to test our hypotheses that structural factors - such as morphometrical asymmetry and connectivity (both inter- and intrahemispheric) - of speech-related cortex were viable substrates for lateralisation of language processing and acuity in rapid temporal processing (i.e., processing acoustic speech-related cues in non-linguistic stimuli). The hypotheses and findings for this set of aims is discussed above in the core body of the text. A secondary set of aims is reported here in the supplementary material. The aims of this material were centred on elucidating the relevance of granularity and nuance in defining a.) ROI for structural measures as well as b.) the measurements themselves (i.e., variables of interest).

Whilst critique of continued use of the definitions of Broca’s and Wernicke’s areas for speech-related cortex has been well articulated in review literature [50-52], such definitions and parcellations remain strongly prevalent [e.g., 20, 174, 175, 176]. A relevant contribution can be made by the present study due to a-priori identifying such parcellation concerns and providing empirical validation to the importance of nuance for ROI.

Regarding variables of interest, superiority of use of constituent components of volume (surface area, thickness, and gyrification) was effectively emphasised nearly 10 years ago [28, 60] and appears to have been well adopted since [e.g., 20, 26, 63]. Our aim was to further verify the value of surface area as well as thickness and local gyrification as variables when studying morphometrical asymmetry.

We expected our findings to reinforce the importance of measuring cortical surface-area and thickness in place of volume. Specifically, we hypothesised that the two constituent measures would not show consistent relationships across global and regional measures (i.e., inverse or null correlations would occur). Furthermore, that surface-area would sufficiently predict volume asymmetries for our ROI, as has been previously reported [28, 60]. Regarding morphometrical asymmetry of our ROI, we hypothesised to replicate the morphometrical asymmetries reported by Chiarello et al. [29] and Kong et al. [9] for the PT, HG, and IFG (in particular the IFGop and IFGtr subcomponents). That is, leftward surface-area asymmetry was predicted for the PT, HG, and IFGop. Although Chiarello and colleagues (n = 250) reported a leftward asymmetry for the IFGtr, Kong and colleagues (n = 17,141) reported the inverse. We hypothesised to replicate the latter findings given the larger sample size.

### Methods

All methods are described in the core body of the paper. In this section exploratory analyses were implemented and as such false discovery rate corrections were applied to unplanned multiple comparisons [177].

### Results

#### Regions-of-interest and their asymmetries

Paired samples t-tests were performed to assess general patterns of structural asymmetry. For surface-area measures the data revealed a significant leftward asymmetry for the PT, *t*(60) = 4.39, *p* < .001, HG, *t*(60) = 7.78, *p* < .001, and IFGop, *t*(60) = 4.94, *p* < .001.

No significant asymmetry was evident for IFGtr. These findings are congruent with those of Chiarello et al. [29], who used the same cortical parcellation methods with a larger sample (n = 200) of similar demographics with the exception that they also observed a significant leftward asymmetry of IFGtr surface-area as well. Kong et al. [9] (n= 17,141) also found these patterns of asymmetry extended to the HG and IFGop (PT was not measured) although they observed a rightward asymmetry of IFGtr.

Our results showed no significant asymmetry of cortical thickness for any ROI, which contrasts Chiarello and colleagues who reported significant rightward thickness asymmetries for each of these ROI. The measures of local regions showed significant rightward asymmetry of gyrification for the PT, *t*(59) = -2.71, *p* = .009 replicating that observed by Chiarello and colleagues, but we did not replicate their observed rightward gyrification asymmetry for IFGop or leftward gyrification asymmetry for IFGtr. It should be noted that a non-significant asymmetry index in our data reflects that our overall sample did not show a general leftward or rightward asymmetry for that particular measure and region, though asymmetries may be meaningfully present in sub-groups and individuals. Supplementary Table 1 provides an overview of these findings.

**Table 1.** Cortical Surface-Area, Thickness, Gyrification, & Asymmetry Index for Regions-of-Interest.

| Planum Temporale |  |  |  |  |  |  |  |  |  |  |  |
| --- | --- | --- | --- | --- | --- | --- | --- | --- | --- | --- | --- |
| Surface-area (mm<sup>2</sup>) |  |  |  | Thickness (mm) |  |  |  | Local Gyrification |  |  |  |
| Left Hemisphere |  | Right Hemisphere |  | Left Hemisphere |  | Right Hemisphere |  | Left Hemisphere |  | Right Hemisphere |  |
| M | SD | M | SD | M | SD | M | SD | M | SD | M | SD |
| 667.54 | 109.13 | 596.1 | 100.06 | 2.73 | 0.17 | 2.72 | 0.18 | 3.93 | 0.28 | 4.01 | 0.26 |
| Asymmetry Index |  |  |  | Asymmetry Index |  |  |  | Asymmetry Index |  |  |  |
| **M = 5.62\*\*** (SD = 9.94) |  |  |  | M = 0.12 (SD = 3.99) |  |  |  | **M = -1.03\*** (SD = 2.91) |  |  |  |
| Heschl's Gyrus |  |  |  |  |  |  |  |  |  |  |  |
| Surface-area (mm<sup>2</sup>) |  |  |  | Thickness (mm) |  |  |  | Local Gyrification |  |  |  |
| Left Hemisphere |  | Right Hemisphere |  | Left Hemisphere |  | Right Hemisphere |  | Left Hemisphere |  | Right Hemisphere |  |
| M | SD | M | SD | M | SD | M | SD | M | SD | M | SD |
| 351.89 | 79.07 | 275.62 | 58.8 | 2.72 | 0.21 | 2.73 | 0.2 | 4.95 | 0.38 | 4.97 | 0.33 |
| Asymmetry Index |  |  |  | Asymmetry Index |  |  |  | Asymmetry Index |  |  |  |
| **M = 11.88\*\*** (SD = 10.86) |  |  |  | M = -.13 (SD = 4.02) |  |  |  | M = -0.26 (SD = 2.73) |  |  |  |
| Inferior Frontal Gyrus – Pars Opercularis |  |  |  |  |  |  |  |  |  |  |  |
| Surface-area (mm<sup>2</sup>) |  |  |  | Thickness (mm) |  |  |  | Local Gyrification |  |  |  |

| Left Hemisphere |  | Right Hemisphere |  | Left Hemisphere |  | Right Hemisphere |  | Left Hemisphere |  | Right Hemisphere |  |
| --- | --- | --- | --- | --- | --- | --- | --- | --- | --- | --- | --- |
| M | SD | M | SD | M | SD | M | SD | M | SD | M | SD |
| 1029.38 | 175.82 | 933.93 | 173.72 | 2.92 | 0.15 | 2.92 | 0.16 | 4.71 | 0.37 | 4.77 | 0.32 |
| Asymmetry Index |  |  |  | Asymmetry Index |  |  |  | Asymmetry Index |  |  |  |
| **M = 4.97\*\*** (SD = 7.25) |  |  |  | M = 0.14 (SD = 2.15) |  |  |  | M = -.58 (SD = 2.74) |  |  |  |

| Inferior Frontal Gyrus – Pars Triangularis |  |  |  |  |  |  |  |  |  |  |  |
| --- | --- | --- | --- | --- | --- | --- | --- | --- | --- | --- | --- |
| Surface-area (mm<sup>2</sup>) |  |  |  | Thickness (mm) |  |  |  | Local Gyrification |  |  |  |
| Left Hemisphere |  | Right Hemisphere |  | Left Hemisphere |  | Right Hemisphere |  | Left Hemisphere |  | Right Hemisphere |  |
| M | SD | M | SD | M | SD | M | SD | M | SD | M | SD |
| 818.29 | 130.53 | 776.62 | 160.64 | 2.86 | 0.17 | 2.83 | 0.17 | 3.88 | 0.28 | 3.87 | .26 |
| Asymmetry Index |  |  |  | Asymmetry Index |  |  |  | Asymmetry Index |  |  |  |
| M = 2.96 (SD = 11.69) |  |  |  | M = 0.47 (SD = 3.13) |  |  |  | M = 0.02 (SD = 3.4) |  |  |  |

*Note.* M = mean, SD = standard deviation, AI = asymmetry index. Positive asymmetry index = leftward asymmetry. \* = significant relationship ( $p < .05$ ), \*\* = significant relationship ( $p < .005$ ).

#### Relation of cortical volume asymmetry to surface-area, thickness, gyrification asymmetry, and total brain volume

A linear stepwise regression of PT volume asymmetry was conducted with surface-area, thickness and gyrification asymmetry, and total brain volume as predictor variables. The preferred model included surface-area only for optimally explaining variance of PT volume asymmetry, *R^2Ad^*^j^ = .819, with inclusion of thickness and gyrification offering only small increases in explained variance (surface-area + thickness, *R^2Ad^*^j^ = .881; surface-area + thickness + gyrification, *R^2Ad^*^j^ = .891). Thus, affirming previous reports, surface-area explains most of the variance of regional volume asymmetry for the PT [29, 60, 178].

#### Exclusion of the inferior frontal gyrus pars triangularis

Our findings did not show significant surface-area asymmetry for the IFGtr. This replicates the findings of Greve et al. [28]. Indeed, recent findings suggest that the IFGop, but not IFGtr is directly connected to perisylvian regions via the arcuate fasciculus [71, 179]. Taken together with previous findings implicating functional links of the PT, HG, and IFGop (but not IFGtr) in rapid temporal and phonetic processing [108, 113, 180-182], this region was excluded from further analyses in our study.

#### Interrelations of measures

Paired samples t-tests affirmed the findings of [9; n = 17,141] that in general the left hemisphere compared to the right was larger in cortical thickness was *t*(60) = 4.21, *p* < .001, and smaller in surface-area, *t*(60) = -2.98, *p* = .004. Local gyrification did not significantly differ between the two hemispheres. Partial bivariate correlations were conducted between the total surface-area, mean thickness, and total gyrification of each hemisphere (see Supplementary Table 2). Consistent with these previous studies, at the hemispheric level surface-area was inversely related to thickness and positively related to local gyrification. Thickness and gyrification did not show any relationship at the hemispheric level. Thus, brain hemispheres with greater surface-area were typically thinner and more folded. Although correlations were significant for both hemispheres, they were notably higher in the right. These findings are again supportive of independence or inverse relationships between the constituent measures of volume.

**Table 2.**
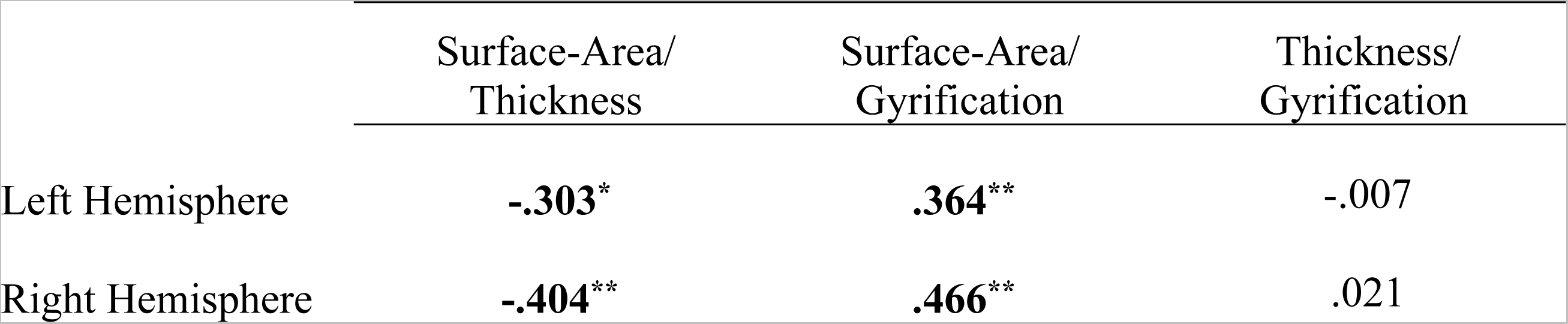
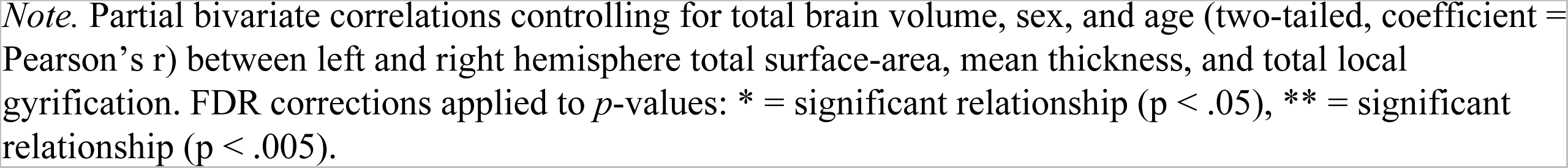
Hemispheric Relationships between Morphometry Measures.

*Hemispheric Relationships between Morphometry Measures*
|  | Surface-Area/<br>Thickness | Surface-Area/<br>Gyrification | Thickness/<br>Gyrification |
| --- | --- | --- | --- |
| Left Hemisphere | **-.303\*** | **.364\*\*** | -.007 |
| Right Hemisphere | **-.404\*\*** | **.466\*\*** | .021 |

*Note.* Partial bivariate correlations controlling for total brain volume, sex, and age (two-tailed, coefficient = Pearson's $r$ ) between left and right hemisphere total surface-area, mean thickness, and total local gyrification. FDR corrections applied to $p$ -values: \* = significant relationship ( $p < .05$ ), \*\* = significant relationship ( $p < .005$ ).

Partial bivariate correlations (two-tailed; controlling for sex and age) were conducted between each measurement asymmetry index of the PT, HG, and IFGop. For the PT, a moderate negative correlation emerged for thickness and gyrification, *r* = −.494, *p* <.001, but no relationship between surface-area and thickness, or surface-area and gyrification was evident. In partial contrast, Chiarello et al. [29] found no between-measure relationships for the PT. For the HG, we replicated their observed inverse relationship between thickness and gyrification, with a weak negative correlation, *r* = −.347, *p* = .008. We did not observe the inverse correlation between surface-area and thickness asymmetry reported by Chiarello and colleagues for the IFGop. Overall, our findings support some of the general patterns reported by Chiarello and colleagues: asymmetry of surface-area and gyrification were not related at the single-region level for PT, HG, or IFGop, whilst thickness and gyrification asymmetry were inversely related only in perisylvian region(s) (both PT & HG in our study, HG only in theirs).

### Structural asymmetries and behavioural abilities

#### Rapid temporal processing with planum temporale and Heschl’s gyrus

As reported in the core paper, partial bivariate correlations (one-tailed; controlling for sex, age, and total brain volume) were conducted to assess for predicted relationships between measures of leftward PT surface-area asymmetry with better RTP ability as measured by GDTs. Similar bivariate correlations (two-tailed) were also used to explore other potential structural asymmetry relationships. As predicted, a weak negative correlation was found for PT surface-area asymmetry, *r* = −.28, *p* = .017. No relationships were evident however between GDTs and PT thickness or gyrification. Likewise, no relationships for any measure of HG or IFGop asymmetry seemed related to GDTs in our sample. Thus, only greater leftward asymmetry of PT surface-area seemed related to behavioural rapid temporal acuity and in the direction of better ability. Although a relationship between leftward asymmetrical PT thickness and behavioural performance on temporal processing tasks has previously been shown using surface-based analyses, the study involved a complex pattern matching task rather than direct RTP measure [69]. A previous study (n = 10) reported that a group with leftward HG volume showed greater leftward lateralisation (measured using functional MRI; functional MRI) to rapid temporal information than a rightward HG volumetric asymmetry group [70]. However, no existing literature is available to compare our results regarding RTP behavioural performance and surface-based morphometric asymmetries of HG.

#### Language lateralisation and planum temporale, Heschl’s gyrus, and inferior frontal gyrus pars opercularis asymmetry

As reported in the core paper, partial bivariate correlations (one-tailed; controlling for sex, age, and total brain volume) were conducted to assess for predicted relationships between leftward PT surface-area asymmetry with stronger leftward language lateralisation as measured by REA in the NF-DLT. Similar bivariate correlations (two-tailed) were used to explore other potential structural asymmetry relationships. As predicted, a weak positive correlation was found for PT surface-area asymmetry, *r* = .245, *p* = .032 (larger surface area asymmetry, better NF-DLT). No relationships were evident however between REA and PT thickness or gyrification asymmetry. Likewise, no relationships for any measure of HG or IFGop asymmetry seemed related to language lateralisation, as measured by NF-DLT, in our sample. Thus, only greater leftward asymmetry of PT surface-area seemed related to behavioural leftward language lateralisation.

As reported in the core paper, partial bivariate correlations (one-tailed; controlling for sex, age, and total brain volume) were conducted to explore the expected relationships of PT-to-PT connectivity with our measures of PT surface-area asymmetry, language lateralisation, and RTP ability. Cortical thickness asymmetry of the PT was also assessed to affirm the specific relevance of surface-area. As predicted, our results showed an inverse correlation of PT-to-PT connectivity and leftward PT surface-area asymmetry, *r* = −.309, *p* = .01, whilst cortical thickness asymmetry of the PT showed no such relationship, *r* = −.099, *p* = .468 (two-tailed).

## Discussion

Our study reinforces that structural asymmetries are a marked feature of the cortex of the human brain [9, 183, 184]. Earlier studies have typically demonstrated this finding using cortical volume as the measure of interest. Our findings align with more recent arguments that such asymmetries are better addressed using the constituent measures of surface-area and thickness, due to their weak genetic links [57, 59] and often absent or inverse morphometric relationships [9] creating nuanced functional relationships to surface area *or* thickness that are absent when measuring both as volume [68]. Indeed, our findings support previous reports that surface-area accounts for a large proportion of variance of volume asymmetries [29, 59, 60, 185]. Including thickness in the regression model offered only a small increase in explained variance. Furthermore, the two asymmetry measures were inversely related at the hemispheric level and generally showed no relationship at the ROI level.

Our findings also highlight that asymmetry patterns are not easily generalisable. That is, patterns at the hemispheric level do not necessarily represent ROI patterns. Although we replicated an inverse relationship of surface-area and thickness, positive relationship of surface-area and gyrification, and no correlation of thickness and gyrification at the hemispheric level, these relationships broke down at the ROI level as previously shown [29]. Surface-area has been positively related to gyrification, and both inversely related to thickness at the regional (multiple ROI) level [58]. However, with the example of the PT, only an inverse relationship for thickness and gyrification emerged in our findings, whilst no such relationships were found by Chiarello et al. [29]. Thus, our findings further the strong support for exploring asymmetry patterns with as much nuance available in terms of both measurement and region outline.

Regarding region outline, our results are affirming of the insufficient precision that the constructs of Broca’s and Wernicke’s areas [50-52]. Our ROI have been merged for the assessment of structural factors of speech and language (PT & HG for Wernicke’s, IFGop & IFGtr for Broca’s) in other studies [20, 174, 176]. As outlined below, such a methodology may however nullify effects observable when regions are more granularly represented, especially in the case of assessing structural asymmetry effects.

### Planum temporale

Our study affirms that leftward surface-area asymmetry of the PT indeed appears to be a general phenomenon in the human brain, and that this constituent measure may best represent prior reports of a volumetric asymmetry. Gyrification asymmetry has been studied only twice (including the present study) yet appears rightward. Cortical thickness asymmetry as a general phenomenon is unclear but is likely unremarkable even if statistically present. Our findings replicated the observed rightward gyrification asymmetry reported by Chiarello and colleagues [29]. We also replicated their lack of correlation between the leftward surface-area and rightward gyrification asymmetry, suggesting that the two effects are independent. In contrast to their findings, we did not replicate a (considerably weak) rightward thickness asymmetry. Meyer and colleagues [60] likewise found no significant thickness asymmetry.

The exploratory analyses of the present study did not show a relationship of PT thickness asymmetry with behavioural RTP ability as measured by GDT, nor language lateralisation as measured by the DLT. This aligns with Qin et al. [27] who found a significant positive relationship of PT surface area, but not thickness with lateralisation of PT activity (measured via fMRI) in a speech comprehension task. Leftward thickness asymmetry has been linked with processing of longer-temporal window cues [i.e., rightward biased under the AST; 69]. Our behavioural tasks were however left-hemisphere biased (RTP and language) under the asymmetric-sampling in time hypothesis [16, 165]. Cortical thickness of the left PT has also been linked to ability for processing of lexical tones where stored templates are available [72], aligning with the computational hub conceptualisation [186]. It should be noted however, that given cortical thickness is only more recently receiving attention, understanding of its relevance is nascent. Indeed, whether thicker or thinner cortex is in general functionally advantageous is yet to be established [60].

Ours is the first study to report local gyrification of the PT in the context of behavioural tasks. Although a general rightward asymmetry was evident, our results showed no significant correlation of such gyrification asymmetry with behavioural RTP ability as measured by GDT, nor language lateralisation as measured by NF-DLT REA. Fuhrmeister and Myers [63] reported surface area and gyrification (but not thickness) of speech-related cortex as factors of differences in phonetic categorisation ability. However, in the case of gyrification this was regarding the bilateral transverse temporale gyri and not measuring asymmetry. Thus, surface area is currently the only measure of the PT with notable relevance for asymmetry as a factor in speech-related processing.

### Heschl’s gyrus

Leftward asymmetry of HG surface-area has been reported as one of the strongest regional brain asymmetries in the general population [9]. Our findings further support this claim. Similar to the PT, cytoarchitectural explanations for asymmetry includes a greater number of microcolumns in the left than right HG. Differing from the PT however, columnar width and spacing asymmetries are not evident [83]. The HG has been implicated as an acoustic substratum of VOT processing, however this is reported as a bilateral sensitivity [181, 187, 188]. Indeed, Chance and others [83] propose that despite apparent structural asymmetries for both primary and associative auditory cortex (i.e., HG and PT), functional lateralisation emerges only in the latter.

Functional lateralisation has been previously reported for HG, though the evidence is not yet compelling. Activation as measured in fMRI response to noise stimuli presented with either increasing temporal or spectral complexity was shown to be greater in the left and right HG respectively in individuals with leftward (but not non-leftward) HG volumetric asymmetry [70]. However, this was a categorical comparison between two groups of only five participants. Greater cortical thickness in the left HG has been associated with superior hearing acuity [68], however this does not reflect asymmetry nor speech-specific auditory processes. Indeed, our results showed no relationship of any HG structural asymmetry measure with behavioural RTP ability as measured by GDT, nor language lateralisation as measured by NF-DLT REA. Thus, the present study indicates that leftward HG surface-asymmetry is a robust effect, yet it does not relate to behavioural language lateralisation or RTP ability. Again, this bears important considerations for merging PT and HG into Wernicke’s area.

### Inferior frontal gyrus pars opercularis

Our findings showed a significant leftward surface-area asymmetry of the IFGop, but no significant surface-area asymmetry of the IFGtr. This pattern has been previously shown using volumetric [73, 189], as well as surface-area measures [28; n = 269]. The leftward asymmetry of IFGop also replicates the studies of Kong et al. [9] and Chiarello et al. [29], though the former found a rightward, and the latter a (considerably weak) leftward surface-area asymmetry of the IFGtr. As mentioned, recent studies show that the IFGtr, unlike the IFGop, is not directly connected to perisylvian regions via the AF [71, 179] or involved in rapid temporal and phonetic processing [108, 113, 180-182]. Thus, our findings further demonstrate dissociation between the two supposed components of “Broca’s area” and emphasises the specificity of leftward asymmetry and RTP to the IFGop in speech lateralisation.

We did not observe a significant asymmetry of the IFGop in terms of cortical thickness or gyrification. Kong et al. [9] likewise found no thickness asymmetry (and did not assess gyrification). Chiarello et al. [29] however, found significant rightward asymmetries for both measures. The occurrence of such inconsistency amongst these studies is unclear, particularly given the consistent analysis methods. Regardless, most pertinent to the present study, structural asymmetry of IFGop was not evidenced as a substrate of RTP ability or language lateralisation. No morphological asymmetry measure of the IFGop in our study showed relationship to RTP or language lateralisation. Thus, although the IFGop has been linked to phonological processing [71, 179] and shows consistent leftward surface-area asymmetry, a functional/behavioural relationship with such structural asymmetry, at least with the suite of tasks employed here, remains unsupported. Functional/behavioural links to structure in general however may be more likely, as thickness of the left IFGop has been linked with ability for pseudoword tone-suffix processing in Swedish speakers [72].

## Conclusion

This study continues understanding of how structural asymmetries of speech-related cortex are associated with behavioural measures of language-lateralisation and rapid temporal processing ability. Morphometrical MRI measures included the surface-area, thickness, and gyrification of left and right PT, HG, IFGop, and IFGtr. We affirmed recent literature that suggests cortical volume is better represented by surface-area due to dissociations with thickness. Furthermore, that each constituent measure (surface-area, thickness, & gyrification) now has supportive literature for a relevance as structural features to speech-related processing. However, that it is surface-area of the PT that seems uniquely relevant in terms of asymmetry. Indeed, asymmetry indexes for each region were described, with the PT, HG, and IFGop showing clear leftward asymmetry, whilst the IFGtr did not. The relationship of leftward PT surface-area asymmetry with behavioural measures was also reported, showed predicted positive associations with both RTP ability (inversely measured using GDTs) and language lateralisation (measured using NF-DLT REA). All other ROI failed to show any such relationship for any measure. We believe that this further supports prior critiques of the use of Broca and Wernicke’s areas as ROI in speech-related processing – especially when assessing asymmetry index as the variable of interest. To reiterate, classical models [1-3] undeniably offered revolutionary early insights into the role of structural asymmetry in speech-related processes. However, with the evolution of the field, we agree that the terms Broca’s area and Wernicke’s area are due for extinction as they misrepresent important subtleties of speech-related cortex and its morphology [50].

In sum, this supplementary material extends our closing point for the core paper, which reports the existence and complexity of relationships of speech-relevant structural asymmetries, functional lateralisation, and behavioural ability. Such complexity exists within both the regions and variables of interest in studies. As such, structural-functional-behavioural effects and relationships may only be observable where appropriate nuance has been methodologically considered and applied to both what and how cortex is measured.

JDB contributed to conceptualization, data curation, formal analysis, project administration, investigation, methodology, visualization, and writing, review, and editing of original draft and final submitted article.

GC contributed to data curation, software, and review and editing of the submitted article. BUF contributed to methodology, formal analysis, and review and editing of the submitted article.

US contributed to methodology, supervision, and review and editing of the submitted article. JT contributed to conceptualization, data curation, formal analysis, project administration, funding acquisition, methodology, resources, supervision, and review and editing of the submitted article.

The authors acknowledge Bryan Paton for guidance in methodology and formal analysis, as well as Alex Provost and Jade D. Frost for assistance in data collection.

